# Reversible CD28 checkpoint modulation by cyclic peptides outperforms biologic blockade under exposure-limited conditions

**DOI:** 10.64898/2026.04.09.717469

**Authors:** Katarzyna Kuncewicz, Saurabh Upadhyay, Yutong Ge, Hongliang Duan, Moustafa T. Gabr

## Abstract

CD28 co-stimulatory blockade is an established therapeutic strategy in autoimmune disease, yet every clinical-stage agent shares a structural limitation: high-affinity, long-lived receptor occupancy that precludes dynamic control of immune suppression. In chronic inflammatory conditions, where prolonged immunosuppression carries infection risk and necessitates treatment interruptions, no existing agent permits rapid restoration of immune function. We report CP8, a disulfide-constrained cyclic peptide antagonist that matches the inhibitory potency of clinical-stage CD28 biologics (FR104, Acazicolcept, and Lulizumab) across primary human immune cells from healthy and ulcerative colitis donors, suppressing IL-2 and IFN-γ production without agonist activity. Unlike these biologics, CP8 enables rapid and near-complete restoration of T-cell function upon compound removal, a property mechanistically inaccessible to antibody-based therapeutics and demonstrated here for the first time for any CD28-targeting agent. In a T-cell transfer colitis model, CP8 maintains efficacy under intermittent dosing and outperforms Acazicolcept, a dual CD28/ICOS inhibitor, under exposure-limited conditions, achieving superior disease suppression, tissue preservation, and cytokine reduction. These results demonstrate that potency and pharmacological persistence are decoupled properties, and reframe cyclic peptides as a superior modality for immune checkpoints where temporal control of signaling is essential to balance efficacy with the risks of chronic immune suppression.

## INTRODUCTION

Immune checkpoint pathways play a central role in controlling the magnitude and persistence of adaptive immune responses and have become critical therapeutic targets in oncology, inflammatory disorders, and transplantation medicine (1–3). The clinical validation of PD-1 and CTLA-4 as pharmacological targets established a principle: discrete nodes of immune signaling can be manipulated to produce outsized therapeutic effects (4–6). Increasing attention is now directed toward costimulatory receptors that regulate the activation threshold of T lymphocytes and shape downstream effector responses (7). Among these receptors, CD28 is a key mediator of T-cell activation through its interaction with the ligands CD80 and CD86 expressed on antigen-presenting cells. Engagement of CD28 amplifies T-cell receptor signaling and promotes cytokine production, proliferation, metabolic reprogramming, and survival of activated lymphocytes (8–10). Dysregulated CD28 signaling has been implicated in multiple inflammatory and autoimmune conditions, including inflammatory bowel disease and rheumatoid arthritis (11–13). Yet no approved or clinical-stage CD28-targeting agent enables reversible, temporally controlled modulation of receptor signaling, a gap with direct consequences for diseases requiring dynamic immune regulation.

Therapeutic approaches targeting immune checkpoints have historically relied on biologic agents such as monoclonal antibodies or receptor fusion proteins capable of engaging extended protein-protein interaction surfaces (14–16). The pharmacological limitations of biologic CD28 modulators are not incidental but are mechanistically inseparable from their mode of action: high-affinity, bivalent receptor engagement combined with Fc-mediated half-life extension produces sustained target occupancy that, by design, resists rapid reversal (17–19). In the context of CD28 in particular, this is consequential; excessive activation of this pathway can trigger severe systemic immune responses, yet the same biologic properties that confer potency also preclude the rapid restoration of T-cell function following cessation of treatment (20). This inability to dynamically control the timing and duration of CD28 blockade defines a key pharmacological gap that no current clinical-stage agent has addressed.

The shallow, solvent-exposed protein-protein interaction interfaces characteristic of immune checkpoint receptors have long resisted small-molecule targeting, a constraint that has reinforced biologic dominance of this target class while leaving the pharmacological limitations of antibody-based agents unaddressed (21–23). Cyclic peptides occupy a distinct chemical space: their constrained conformations enable engagement of extended protein surfaces inaccessible to small molecules, while their synthetic tractability permits precise pharmacological tuning that is structurally unavailable to biologics (24–27). Cyclic peptide scaffolds have successfully targeted challenging protein-protein interaction interfaces across extracellular receptors and intracellular signaling proteins (28,29). Unlike Fc-containing biologics, which undergo neonatal Fc receptor-mediated recycling and sustained tissue distribution, cyclic peptides follow conventional clearance mechanisms, enabling exposure-dependent control of target engagement that is pharmacologically inaccessible to antibody-based therapeutics.

Experimental selection approaches enable exploration of vast peptide sequence spaces and have yielded ligands with high affinity and structural complementarity for protein-protein interaction interfaces that resist conventional drug discovery (30–33). We applied such strategies to the CD28 extracellular domain to identify cyclic peptide modulators with controllable pharmacological properties. In the present study, we identified cyclic peptide ligands that modulate the CD28 immune checkpoint with reversible, exposure-dependent control of receptor signaling, a pharmacological property structurally inaccessible to biologic agents. We reasoned that the rapid clearance kinetics and absence of Fc-mediated recycling inherent to cyclic peptides would permit temporal control of CD28 blockade not achievable with antibody-based therapeutics, and evaluated this hypothesis in physiologically relevant cellular and in vivo systems.

## RESULTS

### Phage display reveals a convergent cyclic peptide binding motif on the CD28 extracellular domain

To engage the featureless extracellular interface of CD28 at the CD80-binding surface (Fig. 1A), we performed phage display selection using a disulfide-constrained cyclic peptide library (Fig. 1B) based on the CX_9_C scaffold, designed to present structured interaction motifs compatible with extended protein-protein interaction surfaces. Following four rounds of biopanning against recombinant human CD28 extracellular domain, we observed progressive enrichment of phage populations, with a marked increase in recovery beginning at round three and further amplification in round four (Fig. 1C). This enrichment profile is consistent with selective expansion of CD28-binding clones rather than nonspecific retention.

**Figure 1.**
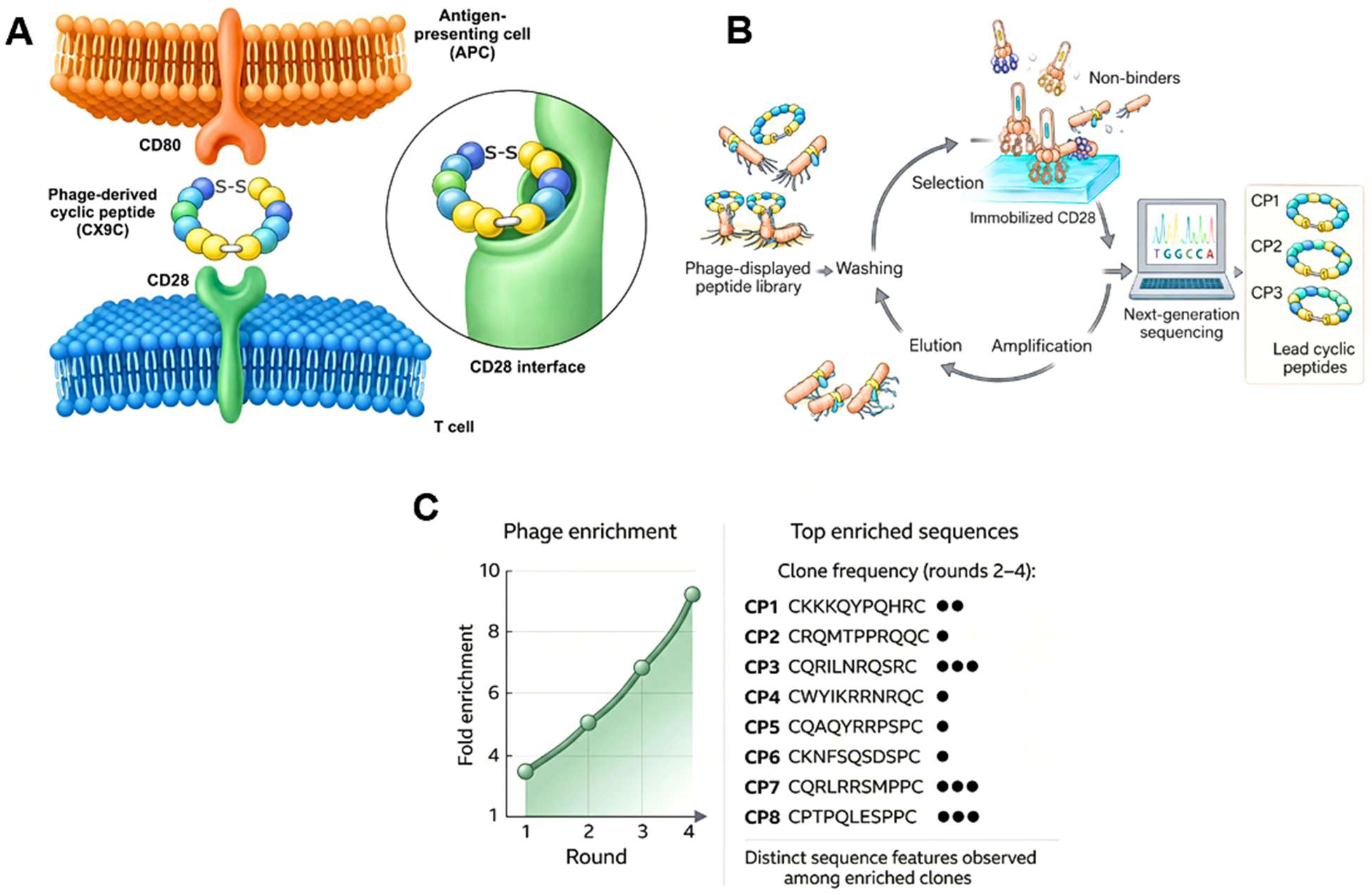
Identification of CD28-binding cyclic peptides by phage display. **(A)** Schematic representation of the targeting strategy. Disulfide-constrained cyclic peptides (CX9C scaffold) were designed to engage the extracellular domain of CD28 on T cells. The approach is motivated by the hypothesis that constrained ligands can access and modulate the CD28 interface involved in receptor-ligand interactions, including the CD28-CD80 axis. The inset shows a conceptual model of peptide engagement at the CD28 surface. **(B)** Phage display selection workflow. A cyclic peptide phage library was subjected to iterative rounds of biopanning against immobilized recombinant human CD28 extracellular domain. Non-binding phages were removed by washing, while bound phage were eluted and amplified for subsequent rounds. After four rounds of selection, enriched pools were analyzed by next-generation sequencing to identify candidate CD28-binding peptides. **(C)** Enrichment and sequence analysis of selected peptides. Progressive enrichment of phage populations across four rounds of selection indicates preferential expansion of CD28-binding clones. Sequencing of enriched pools identified a set of cyclic peptide sequences (CP1-CP8), with relative clone frequencies indicated. The sequences retain the CX9C architecture but exhibit diverse residue compositions, consistent with multiple sequence solutions for CD28 engagement.

Sequence analysis of enriched pools revealed a non-random distribution of cyclic peptide sequences, with multiple clones recurring across independent rounds of selection, as reflected by their relative frequencies (Fig. 1C). While all sequences preserved the CX_9_C architecture, they exhibited diverse residue compositions, indicating that CD28 engagement can arise from multiple sequence solutions rather than a single dominant motif.

Functional interrogation by phage ELISA confirmed that enriched clones exhibited robust and specific binding to immobilized CD28, with signal intensities significantly exceeding background controls (fig. S1). Notably, not all enriched sequences displayed equivalent binding, indicating that sequence enrichment alone does not fully predict functional engagement and highlighting the presence of a functionally privileged subset of binders. Collectively, these results demonstrate that cyclic peptide phage display can uncover ligandable features on the CD28 extracellular domain, despite the absence of a canonical binding pocket.

### Biophysical validation of CD28 binding

Following the synthesis of the peptides (CP1-CP8), their binding to the human CD28 extracellular domain (ECD) was evaluated using microscale thermophoresis (MST) in spectral shift mode. This solution-phase, label-free approach was selected to reliably assess peptide interactions with the relatively flat, protein-protein interaction-driven CD28 surface.

Among the tested peptides, CP1 and CP2 exhibited measurable but weak binding to CD28, with apparent dissociation constants (Kd) of 41.51 µM (95% CI: 22.6-219 μM) and 54.64 µM (95% CI: 38.8-97.3 μM), respectively (fig. S2,3). These interactions were characterized by shallow binding transitions, consistent with low-affinity engagement at an extended protein-protein interface.

In contrast, peptides CP6 and CP8 showed substantially stronger binding behavior, with affinities in the nanomolar range. CP6 bound CD28 with a Kd of 448 nM (95% CI: 310-679 nM), while CP8 displayed further improved affinity, yielding a Kd of 384 nM (95% CI: 270-614 nM) (Fig. 2A,B). The ∼70–150-fold increase in affinity observed for CP6 and CP8 relative to CP1 and CP2 highlights a strong dependence of CD28 engagement on peptide sequence and conformational determinants. The remaining peptides did not exhibit detectable binding to CD28 under the assay conditions. To independently validate the top binder, CP8 was further characterized by isothermal titration calorimetry (ITC), providing direct thermodynamic evidence of interaction (Fig. 2C). Titration of the peptide into CD28 ECD produced clear heat release signals consistent with binding, and fitting to a one-site model yielded a submicromolar affinity (Kd = 2.65 × 10^-7^ M, Fig. 2C). The interaction was predominantly enthalpy-driven, indicative of specific intermolecular contacts such as hydrogen bonding and electrostatic interactions.

**Figure 2.**
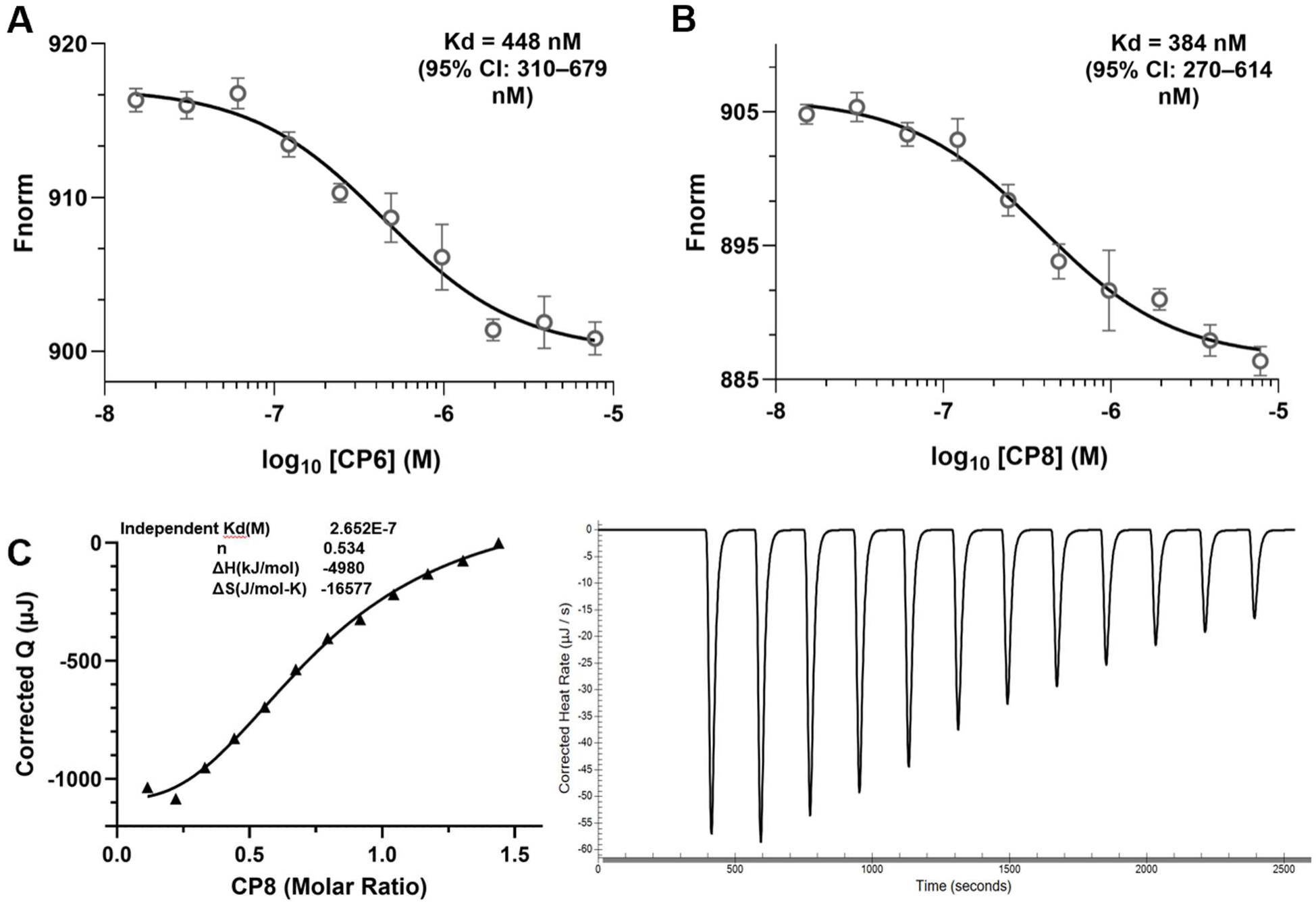
Biophysical analysis of cyclic peptide binding to CD28. Binding the cyclic peptides to the human CD28 extracellular domain (ECD) was quantified by microscale thermophoresis (MST) using intrinsic fluorescence detection (spectral shift mode; Monolith X). Fnorm (670/650 nm) is plotted as a function of peptide concentration. **(A)** CP6 exhibits concentration-dependent CD28 binding. **(B)** CP8 MST binding curve to CD28. Data points represent mean ± SD from three independent measurements. Binding curves were fitted using nonlinear regression, assuming a 1:1 binding model to estimate equilibrium dissociation constants (Kd). **(C)** Thermodynamic characterization of peptide CP8 binding to CD28 by isothermal titration calorimetry (ITC). Representative ITC thermograms (right panel) and the corresponding integrated binding isotherm (left panel) for titration of peptide CP8 into CD28 protein. Heat released upon each injection was integrated and plotted against the molar ratio of peptide to CD28 and fitted using a one-site binding model. Binding analysis yielded a dissociation constant Kd = 2.65 × 10^-7^ M, with an enthalpy change ΔH = −4.98 kcal/mol and binding stoichiometry n = 0.53.

### CP6 and CP8 competitively antagonize CD28-ligand interactions

The ability of top CD28-binding peptides (CP6 and CP8) to functionally disrupt CD28-CD80 interactions was evaluated using an ELISA-based competition assay. Recombinant human CD28 was immobilized on 96-well plates and incubated with biotinylated CD80 in the presence of serial dilutions of peptides. Bound ligand was quantified by streptavidin-HRP detection, and concentration-response curves were derived from three independent experiments. Consistent with the biophysical binding data, both CP6 and CP8 exhibited dose-dependent inhibition of the CD28-CD80 interaction with IC_50_ values in the nanomolar range (Fig. 3A,B). CP6 emerged as the most potent inhibitor with an IC_50_ of 206.10 nM (95% CI: 139-290 nM), while CP8 inhibited CD28-CD80 binding with an IC_50_ of 654.4 nM (95% CI: 430-966 nM).

**Figure 3.**
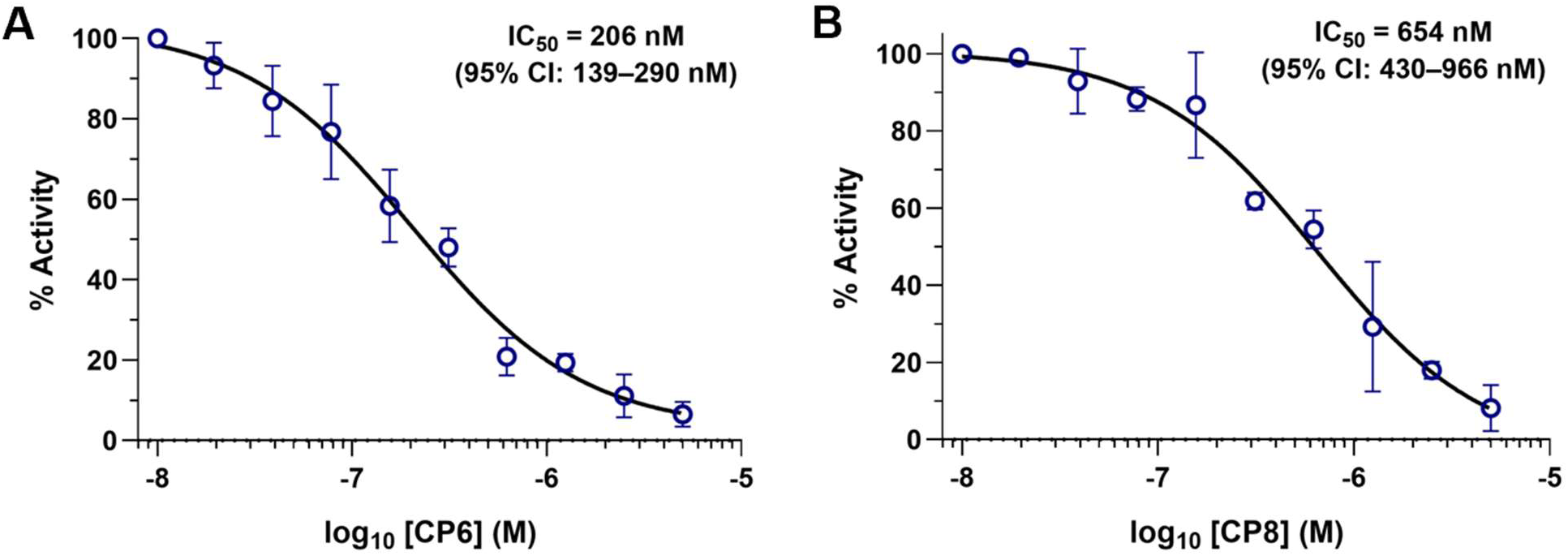
Cyclic peptides competitively disrupt CD28-CD80 interactions. Functional inhibition of CD28-ligand engagement by cyclic peptides was assessed using a competitive binding assay in which peptide-mediated blockade of biotinylated CD80 binding to immobilized recombinant human CD28 was quantified. **(A)** CP6 inhibits CD28-CD80 interaction in a concentration-dependent manner. **(B)** CP8 inhibits CD28-CD80 interaction in a concentration-dependent manner. Data represents SD from three independent experiments. IC_50_ values were determined by nonlinear regression analysis.

### Computational analysis of the CD28 binding mode for CP6 and CP8

To define the structural basis of peptide-mediated CD28 antagonism, we performed molecular dynamics simulations to characterize the binding mode and dynamic stabilization of the CD28-peptide complex. Molecular dynamics (100 ns) simulations of apo CD28 and the CD28-CP6 complex revealed ligand-induced conformational stabilization (Fig. 4A). The complex exhibited an initial equilibration phase (0-50 ns; RMSD ∼1.5 Å), followed by transient conformational rearrangement (50-90 ns; RMSD up to ∼3.5 Å), and convergence to a stable state (90-100 ns; RMSD ∼2.5-3.0 Å), consistent with induced-fit binding. Residue-level analysis (RMSF) showed that regions corresponding to the peptide-binding interface (V5-C22 and F66-T78) were highly flexible in apo CD28 but became markedly stabilized upon peptide binding (Fig. 4B), indicating restriction of local dynamics upon complex formation.

**Figure 4.**
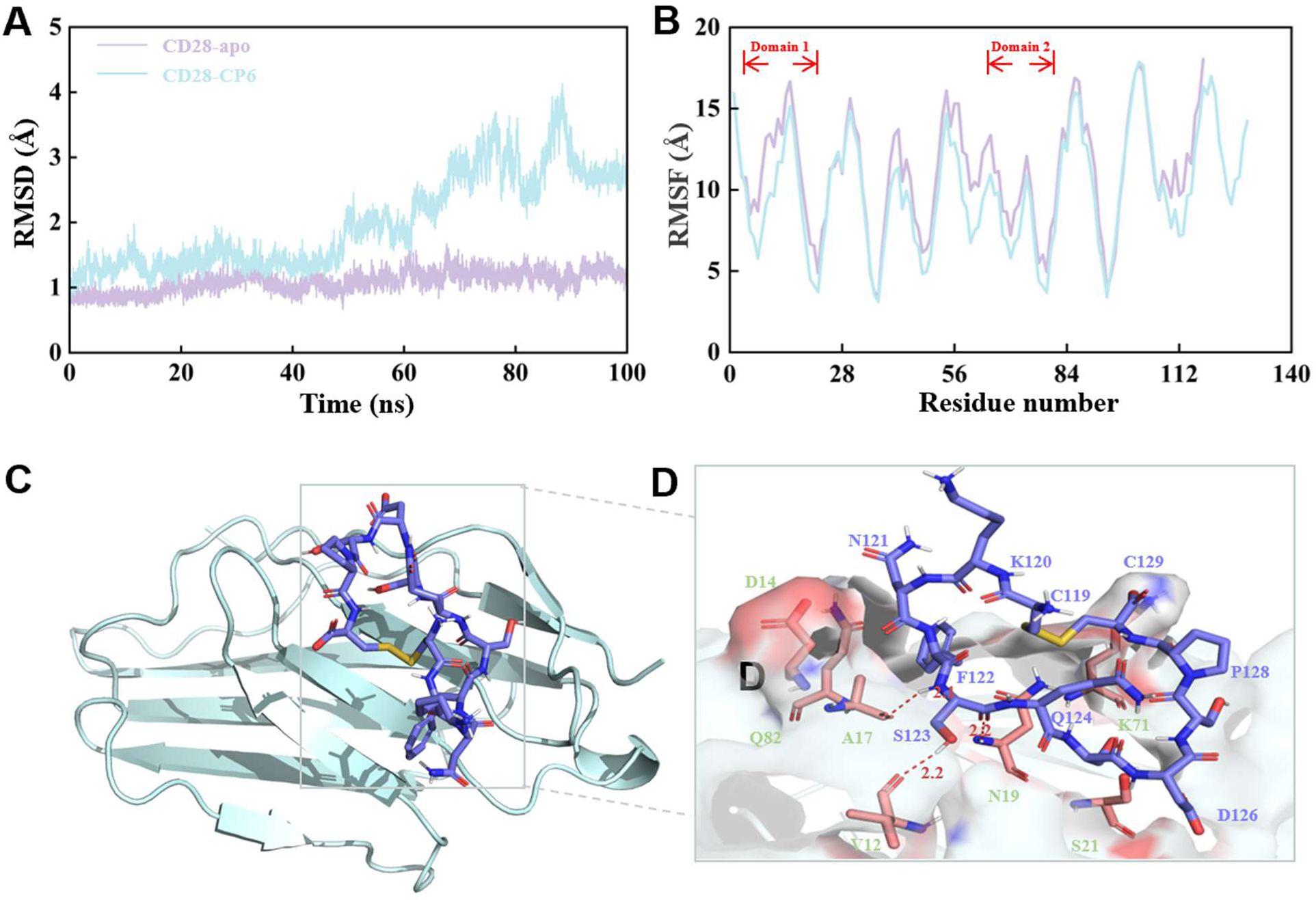
Molecular dynamics evolution and representative binding mode of CD28-CP6. **(A)** Time-dependent root-mean-square deviation (RMSD) curves of the Apo-CD28 receptor (purple line) and the CD28-CP6 complex (light blue line) during the 100 ns molecular dynamics simulation. **(B)** Comparison of the root-mean-square fluctuation (RMSF) of backbone carbon atoms between CD28-apo and CD28-CP6. **(C)** The representative 3D binding conformation of CD28-CP6 extracted via clustering from the stable trajectory at the late stage of the molecular dynamics simulation. The receptor protein CD28 is shown as a blue cartoon model, and the cyclic peptide CP6 is displayed as a purple ball-and-stick model. **(D)** A zoomed-in view of the binding pocket. The receptor surface is represented by a semi-transparent electrostatic potential. Red dashed lines and annotated numbers indicate the key polar hydrogen bond networks and their distances (in Å).

Clustering of the equilibrated trajectory (90-100 ns) identified a dominant binding conformation in which CP6 engages the CD28 surface at a defined interface (Fig. 4C,D). The interaction is anchored by a polar network centered on S123, which forms multiple hydrogen bonds with V12, A17, and N19, while F122 inserts into a hydrophobic pocket, contributing van der Waals stabilization. The intramolecular disulfide bond constrains peptide conformation, enabling precise positioning of key residues. Together, these data define a binding mode in which polar anchoring and hydrophobic packing collectively stabilize the CD28-CP6 complex.

In parallel, we performed 100 ns molecular dynamics simulations of the CD28-CP8 complex to characterize its binding dynamics (Fig. 5A). The trajectory revealed a two-stage induced-fit process, with an initial metastable phase (0-85 ns; RMSD ∼1.5-2.0 Å) followed by a late conformational transition (85-100 ns; RMSD up to ∼3.0 Å), consistent with crossing a secondary energy barrier to access a deeper binding mode. Residue-level analysis showed pronounced stabilization of the CD28 interface upon CP8 binding, with marked reductions in flexibility across Domain 1 (N1-C22) and Domain 2 (V77-T89) (Fig. 5B), indicating restriction of interfacial dynamics following induced-fit rearrangement.

**Figure 5.**
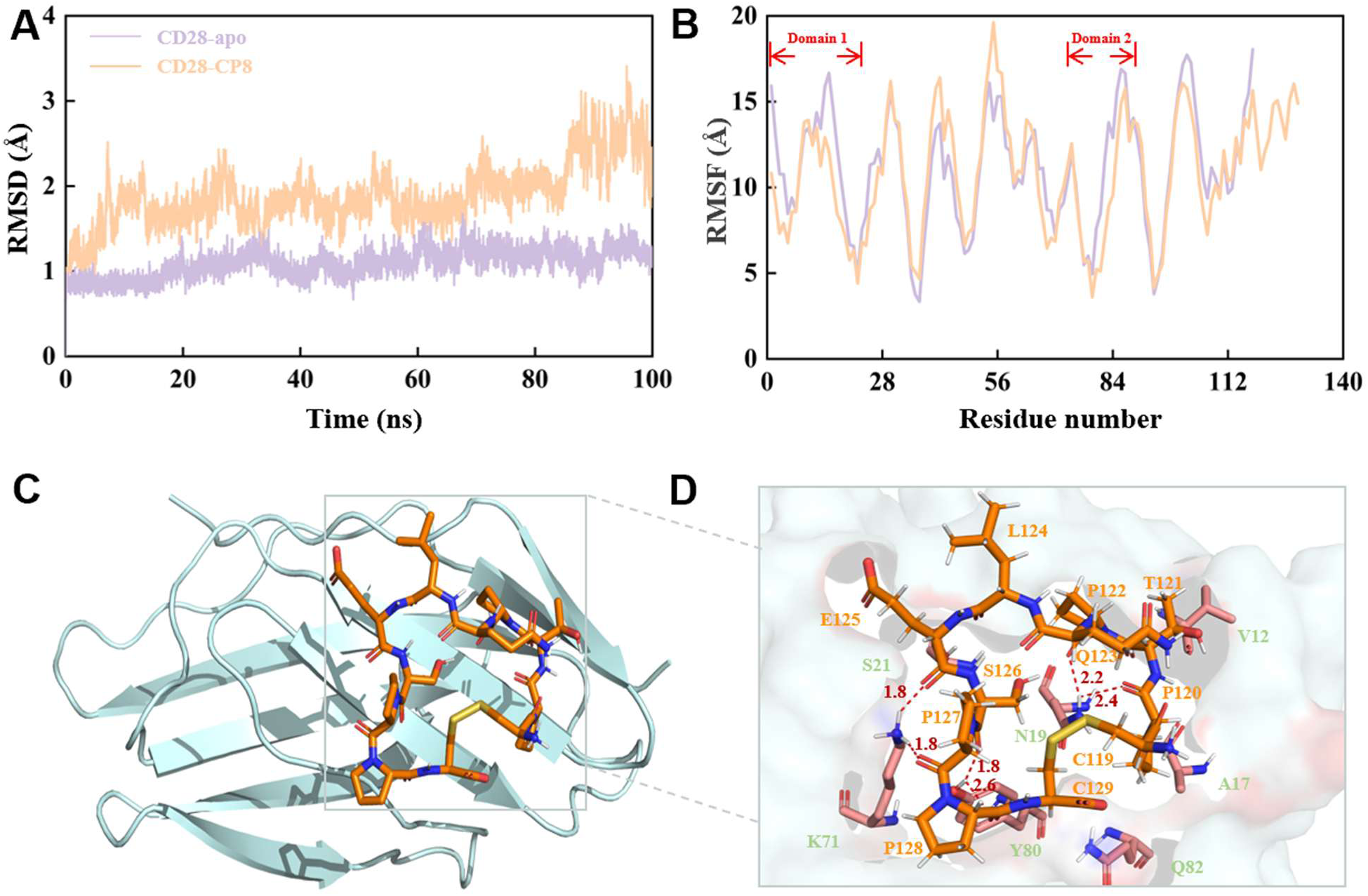
Molecular dynamics simulation and representative binding mode of CD28-CP8. **(A)** Time-dependent root-mean-square deviation (RMSD) curves of the receptor protein CD28 (Apo, purple line) and the CD28-CP8 complex (orange line) during the 100 ns molecular dynamics simulation. **(B)** Comparison of the root-mean-square fluctuation (RMSF) of backbone carbon atoms between CD28-apo and CD28-CP8. **(C)** The representative 3D binding conformation of CD28-CP8 extracted via clustering from the stable trajectory at the late stage of the molecular dynamics simulation. The receptor protein CD28 is shown as a light blue cartoon model, and the cyclic peptide CP8 is displayed as an orange ball-and-stick model. **(D)** A zoomed-in view of the binding pocket. The receptor surface is represented by a semi-transparent electrostatic potential. Red dashed lines and annotated numbers indicate the key polar hydrogen bond networks and their distances (in Å).

Clustering of the equilibrated trajectory identified a dominant binding conformation characterized by a rigid, pre-organized peptide scaffold and an extensive hydrogen-bonding network (Fig. 5C,D). The proline-rich backbone and intramolecular disulfide constraint enforce structural rigidity, enabling precise engagement of key residues. Q123 and P120 interact with N19, while E125 contacts K71, and additional contacts with Y80 further stabilize the complex. Together, these results define a well-defined binding mode in which a conformationally constrained peptide scaffold and multi-point polar interactions collectively stabilize CD28-CP8 engagement.

### Reporter-based functional inhibition of CD28 signaling

To determine whether peptide binding translated into functional inhibition of CD28 costimulatory signaling, the lead candidates (CP6 and CP8) were evaluated using the Promega CD28 Blockade Bioassay, a luciferase-based reporter system that measures CD28-dependent activation in Jurkat effector cells co-cultured with antigen-presenting cells. In this assay, inhibition of CD28 engagement results in reduced reporter activity, providing a quantitative measure of functional CD28 blockade in a cellular environment.

Dose-response experiments were conducted using a 10-point serial dilution series spanning the micromolar to nanomolar range. Both peptides produced clear concentration-dependent inhibition curves, indicating effective suppression of CD28 signaling. The CP6 exhibited moderate inhibitory activity with an IC_50_ of 3.21 μM (fig. S4A), whereas the CP8 demonstrated significantly stronger activity with an IC_50_ of 719.5 nM (fig. S4B). The improved potency observed for CP8 suggests more efficient disruption of CD28-mediated costimulatory signaling at the cellular level. Overall, these results confirm that peptide engagement of the CD28 extracellular domain can translate into measurable functional inhibition in a cell-based assay and validates CP8 as the lead peptide from this work.

### CP8 achieves biologic-level CD28 inhibition while enabling reversible and non-agonistic immune modulation

To determine whether the lead cyclic peptide CP8 can modulate CD28-dependent T-cell activation in physiologically relevant systems, peripheral blood mononuclear cells (PBMCs) from independent healthy donors (n = 10) were stimulated with anti-CD3/CD28 antibodies in the presence of increasing concentrations of CP8. The peptide induced a robust, concentration-dependent reduction in cytokine production, with progressive decreases in IL-2 and IFN-γ observed across all donors (Fig. 6A,B). Despite expected inter-individual variability in baseline responses, the inhibitory effect was consistently reproduced, indicating a stable pharmacological profile across heterogeneous human immune samples.

**Figure 6.**
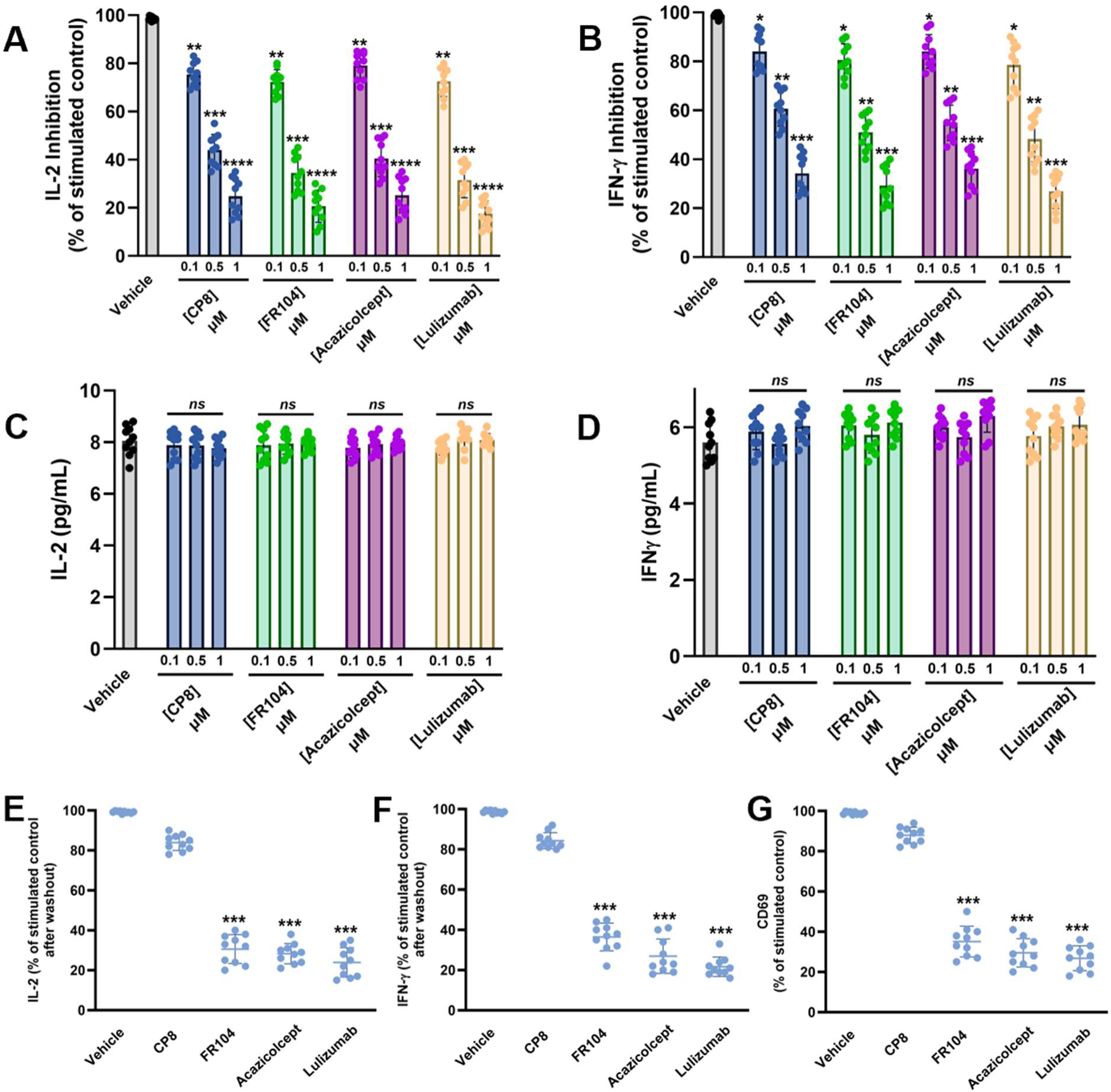
CP8 achieves biologic-level CD28 inhibition while enabling reversible and non-agonistic control of T-cell activation. **(A,B)** CP8 suppresses CD28-mediated cytokine production in primary human PBMCs with efficacy comparable to clinically advanced CD28-targeting biologics. PBMCs from independent donors (n = 10) were stimulated with anti-CD3/CD28 antibodies in the presence of increasing concentrations (0.1, 0.5, 1 µM) of CP8 or the biologic inhibitors FR104, Acazicolcept, and Lulizumab. CP8 reduced **(A)** IL-2 and **(B)** IFN-γ levels in a concentration-dependent manner, reaching levels comparable to biologic-mediated inhibition across donors. **(C,D)** CP8 does not induce cytokine production in the absence of stimulation. Unstimulated PBMCs treated with CP8 or biologic inhibitors across the same concentration range did not exhibit increased **(C)** IL-2 or **(D)** IFN-γ secretion relative to vehicle controls, indicating absence of intrinsic agonist activity. **(E-G)** CP8 enables rapid restoration of T-cell function following inhibitor removal. PBMCs were preincubated with inhibitors, subjected to washout, and restimulated. CP8-treated cells showed robust recovery of **(E)** IL-2 and **(F)** IFN-γ production, as well as **(G)** CD69 expression, approaching stimulated control levels. In contrast, cells treated with biologic inhibitors remained functionally suppressed despite washout, indicating persistent post-exposure inhibition. Data are presented as mean ± SEM with individual donor values shown.

To benchmark functional activity against clinically advanced CD28-targeting modalities, CP8 was evaluated alongside FR104 (Pegrizeprument), Acazicolcept (dual CD28/ICOS inhibitor), and Lulizumab under matched experimental conditions. Across donors, CP8 achieved levels of cytokine suppression comparable to these biologic agents, demonstrating that a compact cyclic scaffold can recapitulate the functional efficacy of established CD28-targeting inhibitors.

Given the historical safety concerns associated with CD28-directed therapeutics, we next assessed whether CP8 induces unintended immune activation in the absence of co-stimulatory input. Treatment of primary PBMCs with CP8 across the tested concentration range did not increase IL-2 or IFN-γ production relative to unstimulated controls (Fig. 6C,D), indicating the absence of intrinsic agonist activity and supporting a favorable baseline functional profile.

We then investigated whether the inhibitory effects of CP8 could be dynamically controlled through reversible receptor engagement. PBMCs were preincubated with CP8 or the indicated biologic inhibitors under conditions yielding comparable initial suppression, followed by removal of unbound ligand and subsequent restimulation. Under these conditions, CP8-treated cells exhibited rapid recovery of cytokine production, reaching approximately 80-90% of stimulated control levels, whereas cells exposed to FR104 (Pegrizeprument), Acazicolcept, or Lulizumab remained markedly suppressed, with recovery limited to ∼20-40% (Fig. 6E,F). Consistent with these findings, CP8-treated PBMCs also restored CD69 expression following washout, whereas biologic-treated cells remained functionally restrained (Fig. 6G).

A defining limitation of biologic CD28 inhibitors is the persistence of functional suppression beyond the exposure window, which constrains temporal control of immune responses. Here, CP8 achieves biologic-level inhibition of CD28 signaling during exposure while, in contrast to these agents, enabling rapid restoration of T-cell activation following removal. These findings demonstrate that CP8 overcomes persistent post-exposure functional restraint, a key pharmacological limitation of current CD28-targeting biologics, thereby converting CD28 blockade from a sustained intervention into a dynamically controllable mode of immune modulation.

### CP8 preserves re-engageable T-cell function in patient-derived immune systems

To determine whether the reversible pharmacology observed in healthy donor systems extends to disease-relevant immune contexts, we evaluated CP8 in PBMCs derived from patients with ulcerative colitis (n = 5). Upon stimulation with anti-CD3/CD28 antibodies, CP8 produced a concentration-dependent reduction in cytokine secretion, attenuating both IL-2 and IFN-γ levels across all patient samples (Fig. 7A,B). The magnitude of inhibition increased with concentration and approached a plateau at 0.5 µM, consistent with effective engagement of the CD28 co-stimulatory pathway under inflammatory conditions.

**Figure 7.**
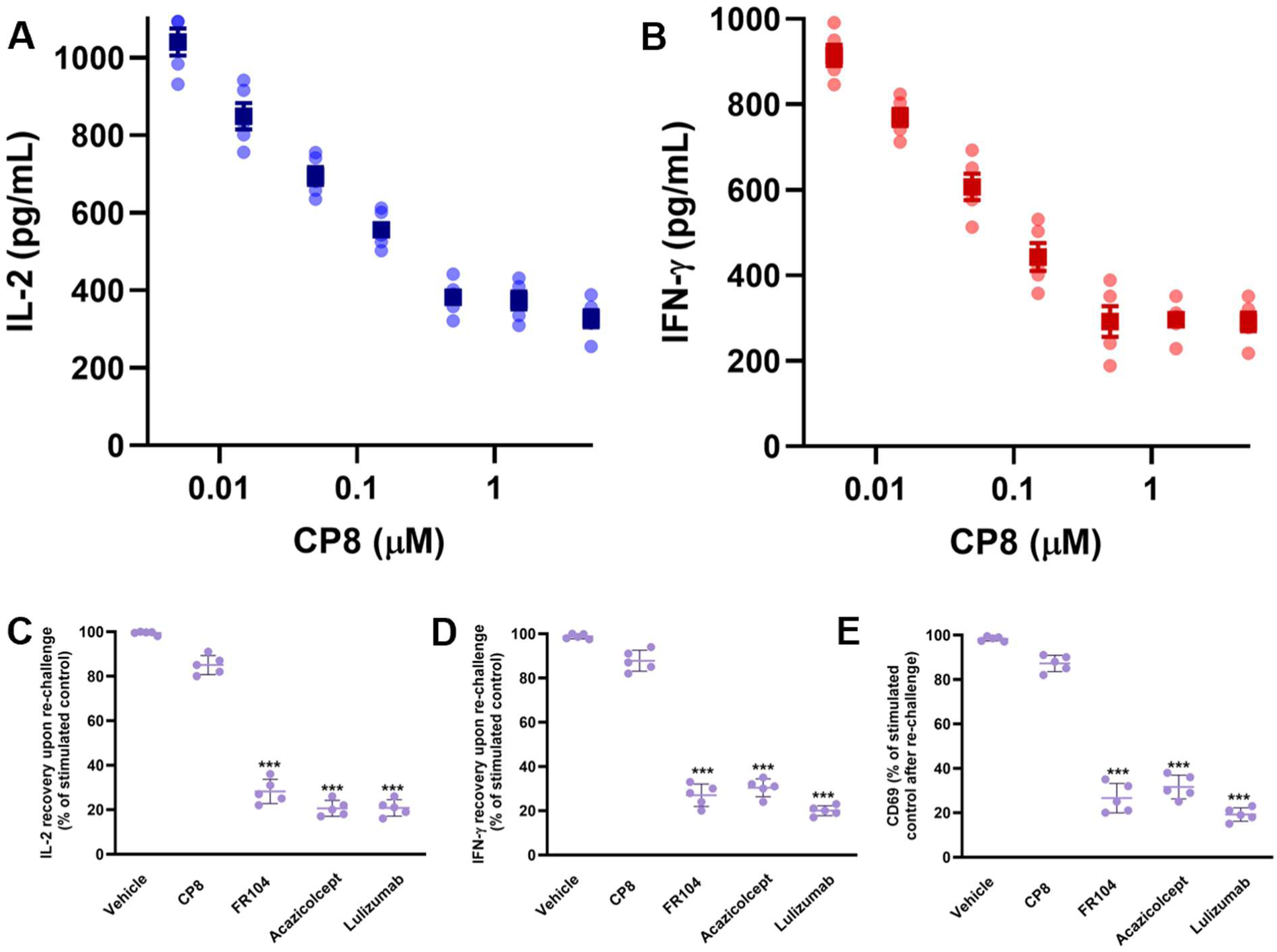
CP8 preserves re-engageable T-cell function in patient-derived immune systems following transient inhibition. **(A,B)** CP8 suppresses cytokine production in peripheral blood mononuclear cells (PBMCs) derived from patients with ulcerative colitis (n = 5). Upon anti-CD3/CD28 stimulation, CP8 reduced **(A)** IL-2 and **(B)** IFN-γ secretion in a concentration-dependent manner, with inhibition approaching a plateau at submicromolar to micromolar concentrations, consistent with effective engagement of the CD28 co-stimulatory pathway under inflammatory conditions. **(C-E)** CP8 preserves the ability of patient-derived T cells to re-engage activation following transient exposure. PBMCs were initially stimulated in the presence of inhibitor, subjected to washout, and subsequently re-stimulated under identical conditions. CP8-treated cells exhibited robust recovery of **(C)** IL-2 and **(D)** IFN-γ production, as well as **(E)** CD69 expression, approaching stimulated control levels upon re-challenge. In contrast, cells exposed to FR104, Acazicolcept, and Lulizumab remained functionally impaired, indicating persistent post-exposure suppression of T-cell responsiveness. Data are presented as individual donor values with mean ± SEM.

To benchmark functional efficacy, CP8 was directly compared with clinically advanced CD28-targeting biologics, including FR104 (Pegrizeprument), Acazicolcept, and Lulizumab, in matched patient samples. Under identical stimulation conditions, CP8 achieved levels of IL-2 suppression comparable to all three biologic agents, confirming preservation of biologic-level inhibitory efficacy in a disease-relevant setting (fig. S5).

We next asked whether CP8 preserves the ability of patient-derived T cells to re-engage activation following transient inhibition. To this end, PBMCs were initially stimulated in the presence of inhibitor, subjected to washout, and subsequently re-stimulated under identical conditions. Following removal of CP8, patient-derived cells exhibited robust recovery of cytokine production upon re-challenge, whereas cells exposed to biologic inhibitors remained markedly impaired (Fig. 7C,D). This effect was consistent across donors and was observed for both cytokine output (Fig. 7C,D) and CD69 activation marker expression (Fig. 7E), indicating restoration of activation competence rather than partial signaling recovery.

Importantly, these findings extend the reversible pharmacology observed in healthy donor systems (Fig. 6) to a disease context, demonstrating that CP8 preserves re-engageable T-cell function in inflammatory immune environments. In contrast, biologic CD28 inhibitors imposed a persistently restrained functional state that limited subsequent activation despite removal of the inhibitor.

Collectively, these results establish that CP8 maintains biologic-level inhibitory efficacy while avoiding prolonged functional suppression in patient-derived immune systems. By enabling T cells to regain responsiveness following transient exposure, CP8 introduces a pharmacological profile characterized by exposure-dependent control of immune activation in disease-relevant human settings.

### CP8 enables exposure-dependent control of disease and outperforms biologic CD28/ICOS blockade under intermittent dosing

To assess the in vivo therapeutic potential of CP8, we employed the adoptive CD4⁺CD45RB^high^ T-cell transfer model of chronic colitis in C.B-17 scid mice, a well-established system that recapitulates key immunopathological features of human inflammatory bowel disease (34–36), including progressive weight loss, elevated disease activity, and sustained inflammatory cytokine production. To ensure translational relevance, we first confirmed that CP8 effectively engages murine CD28. In primary murine splenocytes stimulated with anti-CD3/CD28, CP8 suppressed T-cell activation and cytokine production in a concentration-dependent manner, demonstrating functional cross-reactivity with the murine CD28 signaling axis (fig. S6).

To support in vivo evaluation, the pharmacokinetic (PK) and metabolic stability profile of CP8 was characterized across in vitro and in vivo systems. CP8 exhibited high stability in mouse plasma, with 91% of the parent compound remaining after 6 h at 37 °C. In liver microsomal assays, CP8 displayed low intrinsic clearance in both mouse and human systems (Cl_int_ = 4.4 and 3.1 µL/min/mg, respectively), corresponding to extended microsomal half-lives of 158 min (mouse) and 196 min (human), consistent with favorable metabolic stability across species.

Following subcutaneous administration in mice, CP8 demonstrated rapid absorption and sustained systemic exposure. At 1 mg/kg, CP8 reached a C_max_ of 1.34 ± 0.18 µg/mL with a T_max_ of 0.5 h, while administration at 5 mg/kg resulted in a higher C_max_ of 7.12 ± 0.84 µg/mL under similar absorption kinetics. CP8 exhibited a terminal half-life of 5.8 ± 0.6 h at 1 mg/kg and 6.3 ± 0.5 h at 5 mg/kg, with apparent clearance values of 0.24 and 0.18 L/h/kg, respectively. Systemic exposure increased in a dose-proportional manner, with AUC_0-24h_ values of 10.6 ± 1.4 µg·h/mL at 1 mg/kg and 56.8 ± 6.3 µg·h/mL at 5 mg/kg.

Importantly, plasma concentrations of CP8 remained above the functional threshold required for CD28 inhibition (∼500 nM) for approximately 9 h at 1 mg/kg and 18 h at 5 mg/kg. This sustained target-relevant exposure supports a model in which therapeutic activity is governed by time above the pharmacologically active concentration range rather than continuous receptor occupancy. Together, these findings support a pharmacokinetic-pharmacodynamic relationship in which therapeutic activity tracks with time above the pharmacologically active concentration, providing a mechanistic rationale for efficacy under intermittent dosing without continuous receptor occupancy.

In the T-cell transfer model (Fig. 8A), vehicle-treated animals developed progressive colitis, as evidenced by increasing disease activity index (DAI) scores over the course of the study (Fig. 8B). In contrast, CP8 treatment significantly attenuated disease progression. Daily administration of CP8 (5 mg/kg) produced a marked reduction in clinical disease scores. Notably, an intermittent CP8 dosing regimen (5 mg/kg every other day) yielded a comparable therapeutic outcome, with no significant loss of efficacy relative to daily treatment, indicating that continuous target saturation is not required for disease control (Fig. 8B).

**Figure 8.**
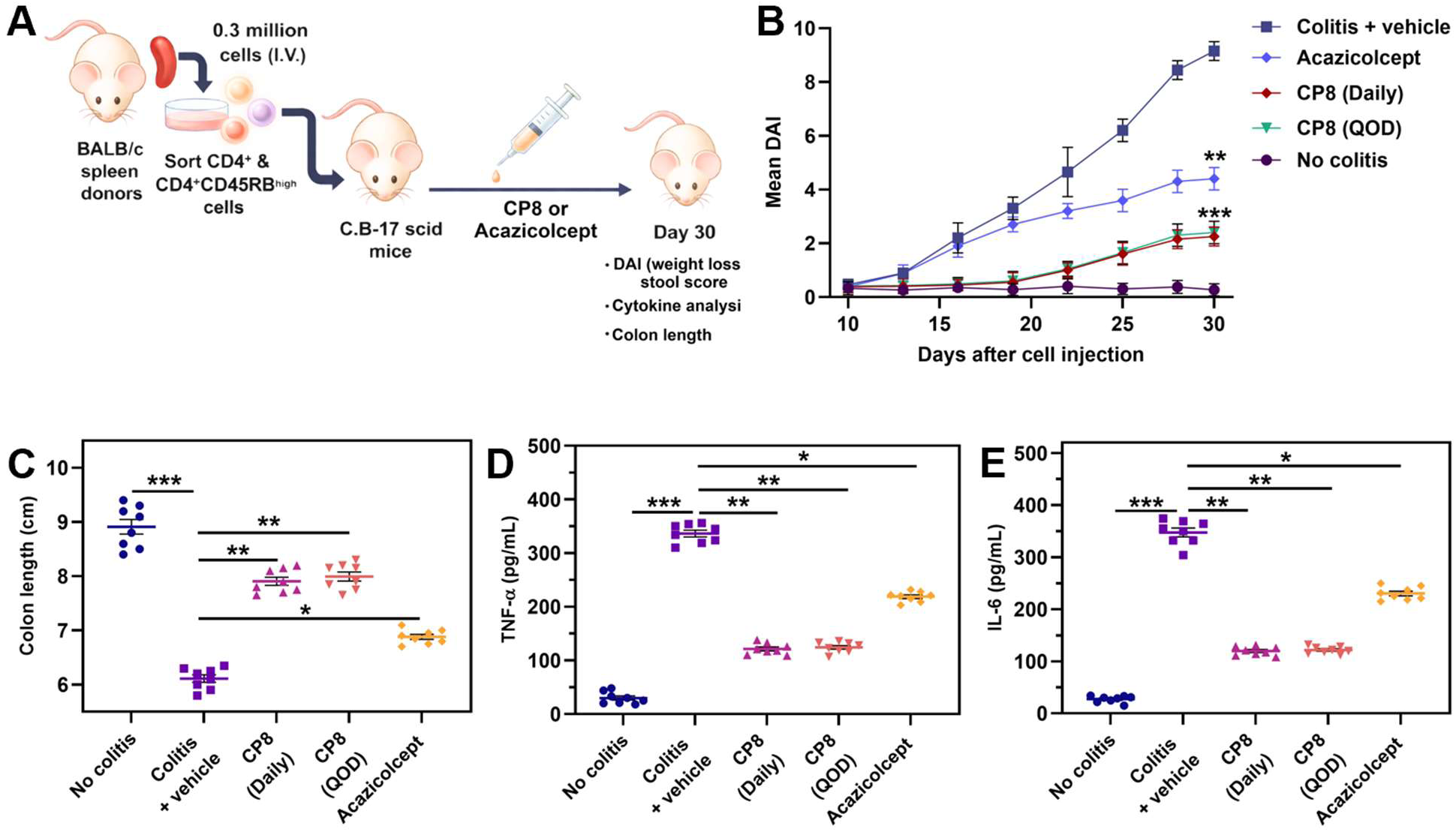
CP8 confers robust and exposure-independent therapeutic efficacy in a T-cell transfer model of chronic colitis and outperforms Acazicolcept. **(A)** Schematic of the adoptive CD4⁺CD45RB^high^ T-cell transfer model. CD4⁺CD45RB^high^ T cells were isolated from BALB/c donor spleens and intravenously transferred into C.B-17 scid recipient mice. Beginning on day 7 post-transfer, mice received CP8 (5 mg/kg; daily or every other day, QOD), Acazicolcept (1mg/kg, twice weekly), or vehicle control until day 30. Disease progression and endpoint analyses included disease activity index (DAI), colon length, and serum cytokines. **(B)** Longitudinal assessment of disease activity index (DAI). Vehicle-treated mice developed progressive colitis, whereas CP8 significantly attenuated disease severity under both daily and QOD dosing regimens. Notably, intermittent CP8 dosing maintained efficacy comparable to daily treatment. Acazicolcept reduced disease severity relative to vehicle but was less effective than CP8 under comparable dosing conditions. Data represent mean ± SEM. **(C)** Colon length at study termination (day 30). Vehicle-treated mice exhibited marked colon shortening, while CP8 treatment (daily and QOD) significantly preserved colon length, indicating reduced tissue inflammation. Acazicolcept treatment resulted in partial restoration relative to CP8. **(D-E)** Serum cytokine levels at endpoint. CP8 significantly reduced systemic inflammatory cytokines TNF-α **(D)** and IL-6 **(E)** under both dosing regimens, with comparable suppression observed between daily and intermittent dosing. Acazicolcept also reduced cytokine levels but to a lesser extent than CP8. Collectively, these data demonstrate that CP8 provides potent disease-modifying activity in a T cell-mediated colitis model and retains efficacy under intermittent dosing, consistent with a reversible and exposure-dependent mechanism of action that distinguishes it from biologic costimulation blockade. Statistical significance was determined using one-way ANOVA with multiple comparisons (\**P* < 0.05, \*\**P* < 0.01, \*\*\**P* < 0.001).

To benchmark CP8 against a clinically relevant costimulation inhibitor, we evaluated Acazicolcept using an established preclinical dosing regimen (1 mg/kg, twice weekly, subcutaneous injection). This dosing regimen was selected to reflect established preclinical schedules for costimulation blockade biologics and to enable comparison under exposure-limited conditions. While Acazicolcept reduced disease severity relative to vehicle, CP8 administered every other day achieved superior control of disease progression, with lower DAI scores across animals. These findings indicate that CP8, a monoselective CD28 inhibitor, maintains therapeutic efficacy under exposure-limited conditions and exceeds the disease-modifying activity of Acazicolcept, a dual CD28/ICOS blocker, when administered at comparable dosing frequencies.

Consistent with clinical outcomes, CP8 treatment preserved colonic architecture. Vehicle-treated mice exhibited pronounced colon shortening at study termination, whereas both daily and intermittent CP8 dosing regimens significantly restored colon length (Fig. 8C), reflecting reduced tissue damage and inflammation. In contrast, Acazicolcept treatment resulted in partial structural rescue, with less complete restoration of colon length relative to CP8-treated groups (Fig. 8C). These data further support the conclusion that transient CD28 modulation by CP8 is sufficient to maintain robust therapeutic benefit.

To further characterize the anti-inflammatory effects of CP8, circulating cytokine levels were measured at the study endpoint. Vehicle-treated animals displayed elevated systemic inflammatory markers, including TNF-α and IL-6 (Fig. 8D,E). CP8 treatment significantly reduced these cytokines under both dosing regimens, with comparable suppression observed in daily and intermittent groups. While Acazicolcept also reduced cytokine levels, the magnitude of suppression was consistently lower than that achieved with CP8, indicating more effective attenuation of pathogenic T-cell-driven inflammation (Fig. 8D,E).

Collectively, these findings demonstrate that CP8 confers robust therapeutic benefit in a T-cell-mediated model of colitis while enabling exposure-dependent control of immune modulation. The ability to maintain efficacy under intermittent dosing establishes that sustained receptor occupancy is not required for disease suppression, distinguishing CP8 from conventional biologic costimulation inhibitors, including Acazicolcept, that rely on prolonged receptor engagement. Importantly, CP8 achieves this as a monoselective CD28 inhibitor, demonstrating that selective CD28 blockade with controlled exposure is sufficient to outperform the pharmacologically persistent dual CD28/ICOS inhibition of Acazicolcept, a finding that implicates the temporal dimension of CD28 signaling control, rather than breadth of target coverage, as the decisive pharmacological variable. Instead, CP8 supports a pharmacological framework in which immune activity can be modulated in a temporally controllable manner, consistent with its reversible and exposure-dependent mechanism of action.

## DISCUSSION

This study establishes cyclic peptides as a therapeutic modality capable of matching the inhibitory potency of biologic CD28 blockade while delivering a pharmacological profile that is structurally inaccessible to antibody-based agents: reversible, exposure-dependent, and temporally controllable. Therapeutic targeting of CD28 has historically relied on biologic agents that achieve potent pathway blockade but remain intrinsically coupled to prolonged receptor engagement and sustained functional suppression. In contrast, CP8 decouples inhibitory efficacy from persistent pathway restraint, demonstrating that effective modulation of CD28 signaling can be achieved through transient and controllable ligand engagement.

A central finding of this work is that CP8 preserves the capacity of T cells to re-engage activation following inhibitor removal. Across both healthy donor and patient-derived immune systems, CP8 achieved levels of cytokine suppression comparable to clinically advanced CD28-targeting biologics, including FR104, Acazicolcept (dual CD28/ICOS inhibitor), and Lulizumab, yet enabled rapid restoration of T-cell function upon washout. In contrast, biologic inhibitors imposed a persistent state of functional restraint despite removal of the agent. These findings reveal a key limitation of existing CD28-targeting strategies, in which pathway inhibition is inherently linked to prolonged suppression of immune responsiveness. By enabling recovery of activation competence following transient exposure, CP8 introduces a mode of checkpoint regulation in which immune activity can be dynamically modulated rather than durably constrained.

Importantly, this reversible pharmacology is preserved in disease-relevant immune contexts. In PBMCs derived from patients with ulcerative colitis, CP8 maintained biologic-level inhibitory efficacy while enabling re-engageable T-cell activation following transient exposure. This demonstrates that the controllable behavior of CP8 extends beyond healthy donor systems to inflammatory immune environments characterized by dysregulated co-stimulatory signaling. The ability to suppress pathogenic activation while preserving the potential for functional recovery represents a critical pharmacological distinction with direct therapeutic implications.

The translational relevance of this mechanism is further supported by in vivo efficacy in the adoptive T-cell transfer model of chronic colitis. CP8 produced robust suppression of disease progression, preserved colonic architecture, and reduced systemic inflammatory cytokines. Notably, intermittent dosing achieved therapeutic outcomes comparable to daily administration, demonstrating that sustained receptor occupancy is not required for disease control. CP8 achieved superior disease control relative to Acazicolcept, indicating that transient modulation of CD28 signaling can outperform biologic costimulation blockade in exposure-limited settings. These findings establish that pharmacological efficacy can be maintained within a defined exposure window, rather than requiring continuous receptor engagement. Critically, this superiority was achieved by a monoselective CD28 inhibitor against a dual CD28/ICOS blocker, with the implication that the therapeutic insufficiency of Acazicolcept under intermittent dosing reflects a pharmacological liability of the biologic modality rather than inadequate target coverage. Temporal control of CD28 signaling alone, when pharmacologically accessible, is sufficient for disease suppression, a conclusion that reframes the rationale for dual blockade strategies and positions controllable CD28 modulation as a distinct and superior therapeutic approach.

Integration of pharmacokinetic and functional data provides a mechanistic framework linking systemic exposure to immune modulation. CP8 exhibits sustained plasma concentrations above the functional threshold required for CD28 inhibition while maintaining a finite exposure window governed by its elimination profile. Together with the rapid restoration of function following washout, these results support a model in which therapeutic activity is driven by time above an effective concentration threshold rather than persistent receptor occupancy. This exposure-dependent mode of action enables predictable and tunable control of immune signaling intensity and duration.

More broadly, these findings highlight a fundamental distinction between synthetic ligands and biologic therapeutics in the context of immune checkpoint modulation. Biologic agents, by virtue of their long half-life and high-affinity binding, are intrinsically optimized for sustained pathway suppression. In contrast, cyclic peptides such as CP8 introduce a pharmacological framework in which efficacy can be temporally controlled, allowing immune responses to be modulated with greater precision. This distinction may be particularly relevant in settings where excessive or prolonged immune suppression is undesirable, including chronic inflammatory diseases and conditions requiring repeated immune engagement.

In conclusion, this work demonstrates that cyclic peptides represent a distinct therapeutic modality, achieving potent inhibition of challenging protein-protein interactions while enabling a pharmacological profile that is structurally inaccessible to biologic agents. By decoupling efficacy from persistence, CP8 redefines the principles of CD28 checkpoint modulation and establishes a framework for next-generation immune modulators based on reversible and exposure-dependent control of signaling pathways.

## MATERIALS AND METHODS

### Phage display selection of CD28-binding cyclic peptides

A disulfide-constrained phage-displayed cyclic peptide library based on a CX_9_C scaffold was used to identify ligands capable of engaging the extracellular domain of human CD28. Recombinant human CD28 extracellular domain (ECD) was immobilized on a solid support, and the phage library was subjected to iterative rounds of biopanning against the target protein. After each round, nonbinding phages were removed by washing, whereas bound phages were recovered by elution and amplified in E. coli for use in the subsequent round of selection. A total of four rounds of biopanning were performed, with enrichment of CD28-binding phage monitored across rounds.

Following the final selection round, phage pools were analyzed by next-generation sequencing to identify enriched peptide sequences. Sequence analysis revealed recurrent CX_9_C-containing clones that increased in frequency over the course of selection, consistent with preferential enrichment of CD28-binding peptides. Representative enriched clones were designated (CP1-CP8) and prioritized for downstream synthesis and functional characterization.

To assess clone-level binding, selected phage were evaluated by phage ELISA against immobilized recombinant CD28 ECD. Briefly, target-coated wells were incubated with individual phage clones, washed to remove unbound material, and bound phage were detected using an anti-M13 horseradish peroxidase-conjugated antibody and colorimetric substrate. Signal intensities were compared with background controls to identify clones exhibiting specific binding to CD28. Clones showing reproducible enrichment and positive phage ELISA signals were advanced for chemical synthesis and orthogonal biophysical validation.

### Peptide synthesis

Peptide synthesis was carried out by solid phase peptide synthesis (SPPS) via the Fmoc/tBu strategy using an automated microwave peptide synthesizer - Liberty BLUE (CEM, Matthews, NC, USA). 424 mg Rink Amide ProTide Resin (0.59 mmol/g; CEM, Matthews, NC, USA) was used to synthesize each peptide. During the synthesis, each of the amino acids was coupled twice, using a 5-fold excess of the amount of resin deposition. The following solutions were used during the synthesis: OxymaPure/DIC as coupling reagents, 20% piperidine in DMF for Fmoc-deprotection, and DMF to wash between the deprotection and coupling steps. Synthesized peptides were cleaved from the resin using a mixture containing: 88% TFA, 5% H2O, 5% phenol and 2% TIPSI (10 ml of solution was used for 424 mg of resin). Crude peptides were precipitated with cold diethyl ether, decanted, and lyophilized. For purification, the peptides were dissolved in water with the addition of a 10-fold molar excess of dithiothreitol (DTT) over free sulfhydryl groups and incubated for 30 minutes at 60 °C. Peptides were purified by reversed phase HPLC, XBridge Prep C18 column (19 x 150 mm, 5 μm, Waters, MA, USA), flow rate 20 mL/min, 10 min run, 5-60% acetonitrile in water with 0.1% formic acid. The purity and mass spectra of the final products were analyzed using an ACQUITY HPLC system equipped with an SQ Detector 2 and an XBridge Prep C18 analytical column (4.6 × 150 mm, 5 μm, Waters, MA, USA), employing a linear gradient from 5% to 100% acetonitrile in water containing 0.1% formic acid over 10 minutes. Oxidation of the peptides was performed using compressed air. The peptide was dissolved in H2O and methanol (1:9, v:v), at a concentration of about 40 mg/L, and the pH was adjusted and kept between 8 and 9 using ammonia. The solution was stirred at room temperature for 7 days, and compressed air was run through the solution. After this time, the solvents were evaporated, and the peptides were lyophilized. Reaction progress was checked using analytical ACQUITY HPLC system equipped with an SQ Detector 2. After this process, the peptides were purified again using the same ACQUITY HPLC system on a XBridge Prep C18 column (19 x 150 mm, 5 μm, Waters, MA, USA). A linear gradient 5%-60% acetonitrile in water with 0.1% formic acid over 10 min was used.

### Recombinant proteins

Recombinant human CD28 extracellular domain (ECD) was obtained from Acro Biosystems (Catalog# CD8-H52H3) and reconstituted according to manufacturer instructions. Proteins were stored in aliquots at −80 °C to avoid repeated freeze-thaw cycles.

### Microscale Thermophoresis (MST)

Binding affinities of CD28-targeting peptides to the human CD28 extracellular domain (ECD) were determined using MST in spectral shift mode. Recombinant His-tagged human CD28 ECD (Acro Biosystems) was fluorescently labeled with RED-tris-NTA 2nd Generation dye (NanoTemper Technologies) according to the manufacturer’s instructions. Labeled protein was diluted to a final concentration of 50 nM in assay buffer consisting of phosphate-buffered saline (PBS, pH 7.0) supplemented with 0.005% (v/v) Tween-20.

Peptides were prepared as serial dilutions in assay buffer and mixed 1:1 with labeled CD28 protein. Samples were incubated for 15 min at room temperature in the dark prior to measurement. MST measurements were performed at 25 °C using a Monolith X instrument (NanoTemper Technologies), and samples were loaded into standard Monolith capillaries.

Fluorescence was recorded at 670 nm and 650 nm, and normalized fluorescence (F_norm) was calculated as the ratio of F670/F650. Dissociation constants (Kd) were obtained by fitting concentration–response curves using a four-parameter nonlinear regression model implemented in MO.Affinity Analysis software and GraphPad Prism 10. Reported values represent mean ± SEM from at least three independent experiments.

### Isothermal Titration Calorimetry (ITC)

Thermodynamic characterization of peptide binding to the CD28 extracellular domain was performed using ITC. Experiments were conducted at 25 °C using a NanoITC calorimeter (TA Instruments). Recombinant human CD28 protein was prepared in PBS buffer (pH 7.4) and loaded into the calorimetric cell at a final concentration of 15 µM.

Peptide solutions were prepared in the same buffer and loaded into the injection syringe at concentrations in the micromolar range. A series of sequential injections (3 µL each) were titrated into the CD28 solution under constant stirring, with sufficient spacing between injections to allow the signal to return to baseline.

Heat released upon each injection was recorded as a function of time to generate raw thermograms. The integrated heat values for each injection were plotted against the molar ratio of peptide to CD28 to generate binding isotherms. Thermodynamic parameters, including the dissociation constant (Kd), enthalpy change (ΔH), entropy change (ΔS), and binding stoichiometry (n), were obtained by fitting the data to a one-site binding model using NanoAnalyze software (TA Instruments).

### Evaluation of CD28-CD80 Binding Inhibition by Competitive ELISA

Inhibition of the CD28-CD80 interaction was evaluated using a competitive ELISA based on the CD28:B7-1 [Biotinylated] Inhibitor Screening Kit (BPS Bioscience, Cat. No. 72007), following the manufacturer’s protocol with minor modifications. Briefly, 96-well plates were coated with recombinant human CD28 (2 µg/mL in PBS; 50 µL per well) and incubated overnight at 4 °C. Plates were washed with Immuno Buffer and blocked with Blocking Buffer for 1 h at room temperature.

Serial dilutions of peptides were added to CD28-coated wells and incubated for 1 h at room temperature to allow binding. Biotinylated CD80 (5 ng/µL) was then added to the wells and incubated for an additional 1 h to assess inhibition of ligand binding to peptide-occupied CD28. Wells lacking CD28 coating served as ligand controls, while wells treated with inhibitor buffer alone were used as negative controls.

After washing, plates were incubated with streptavidin-HRP (1:1000 dilution in Blocking Buffer) for 1 h at room temperature. Chemiluminescent substrate was added, and luminescence was immediately measured using an Infinite M1000 Pro microplate reader (Tecan, Switzerland). IC_50_ values were determined by fitting concentration-response curves using a four-parameter logistic regression model in GraphPad Prism 10. All experiments were performed in triplicate.

### Computational analysis of CD28 binding mode

In this study, molecular dynamics (MD) simulations were performed at 300 K for three systems: the apo CD28 receptor protein (CD28-apo), the CD28-CP6 complex, and the CD28-CP8 complex, using the AMBER20 software package (37) combined with the ff14SB force field (38). All solutes were placed in a truncated octahedral water box and solvated using the TIP3P water model (39), with the minimum distance from the solute surface to the box boundary set to 15.0 Å. Prior to the formal MD simulations, a two-stage energy minimization process was carried out on the systems to eliminate unfavorable steric clashes: (1) first, under positional constraints applied to the solutes, 5000 steps of steepest descent (SD) and 5000 steps of conjugate gradient (CG) energy optimization were performed; (2) subsequently, all constraints were removed, and an unconstrained energy minimization consisting of another 5000 steps of SD and 5000 steps of CG was executed.

After energy optimization, the systems underwent a heating process under positional constraints, during which the temperature was gradually increased from 0 K to 300 K. Subsequently, a 100 ns production simulation under the NPT (constant number of particles, pressure, and temperature) ensemble was conducted at 300 K and 1.0 bar. During the simulation, the temperature was controlled using the Nose-Hoover chain method (40), and the pressure was regulated by the Berendsen method (41). The SHAKE algorithm (42) was employed to constrain all covalent bonds involving hydrogen atoms. The cutoff radius for non-bonded interactions was set to 10.0 Å, and long-range electrostatic interactions were calculated using the smooth Particle Mesh Ewald (PME) algorithm (43). The integration time step was set to 2 fs, and trajectory conformations were saved every 10 ps. Consequently, a total of 10,000 conformational frames were collected for each system for subsequent structural and interaction analyses.

### Cell-Based CD28 Signaling Reporter Assay

Functional inhibition of CD28-mediated costimulatory signaling was evaluated using the CD28 Blockade Bioassay (Promega, Cat. #JA6101). Jurkat CD28 Effector Cells (2 × 10^4^ cells per well) were seeded in white 96-well plates and pre-incubated with serial dilutions of cyclic peptides (10-point, 1:1 dilution series starting at 200 μM, final 1% DMSO) for 1 h at room temperature. An anti-CD28 control antibody (Promega, Cat. #K1231) was included as a positive control.

Following peptide pre-incubation, aAPC/Raji cells (2 × 10^4^ cells per well) were added, and co-cultures were incubated for 5 h at 37 °C in a humidified atmosphere containing 5% CO_2_. Luminescence was subsequently measured according to the CD28 Blockade Bioassay protocol using a GloMax^®^ Discover System (Promega).

Dose–response curves were generated using GraphPad Prism 10 by fitting the data to a four-parameter logistic regression model to determine IC_50_ values. All experiments were performed in triplicate, and results are reported as mean ± SEM.

### Primary human PBMC isolation and culture

PBMCs from healthy donors were obtained from STEMCELL Technologies (Catalog# 70025) and cultured according to manufacturer recommendations. Cells were maintained in RPMI 1640 medium supplemented with 10% fetal bovine serum (FBS), L-glutamine, and penicillin-streptomycin (100 U/mL and 100 µg/mL, respectively) at 37 °C in a humidified atmosphere containing 5% CO_2_. Cell viability following thawing was assessed using trypan blue exclusion prior to experiments.

### T-cell activation assays in primary PBMCs

PBMCs from healthy donors (n = 10 independent donors; STEMCELL Technologies, Catalog# 70025) were cultured in RPMI 1640 supplemented with 10% FBS and antibiotics. Cells were seeded at 2 × 10^5^ cells per well in 96-well plates. T-cell activation was induced using anti-human CD3 (plate-bound, 2 µg/mL) and soluble anti-human CD28 (1 µg/mL). CP8 was added at indicated concentrations (0.005-5 µM) at the time of stimulation. FR104 (MedChemExpress, Catalog# HY-P990587), Acazicolcept (MedChemExpress, Catalog# HY-P99420), or Lulizumab (MedChemExpress, Catalog# HY-P99302) were included as a benchmark controls. Supernatants were collected after 24 h, and IL-2 and IFN-γ concentrations were quantified using ELISA kits (STEMCELL Technologies, Catalog# 02006 and 02003). Each donor sample was tested in technical triplicate.

### Agonist activity assessment

To evaluate intrinsic agonist potential, PBMCs were incubated with cyclic peptides in the absence of stimulation. Cytokine production was measured as described above and compared with baseline controls. Each donor sample was tested in technical triplicate.

### Washout and reversibility experiments

To assess the reversibility of CD28 inhibition, primary human PBMCs were preincubated with 500 nM of CP8, FR104 (MedChemExpress, Catalog# HY-P990587), Acazicolcept (MedChemExpress, Catalog# HY-P99420), or Lulizumab (MedChemExpress, Catalog# HY-P99302) for 60 min at 37 °C prior to stimulation. Following pretreatment, cells were washed three times with pre-warmed RPMI 1640 medium, with centrifugation at 400 × g for 5 min between washes, to remove unbound compound or antibody. Cells were then immediately restimulated with anti-CD3/CD28 activation reagents under the same conditions used for stimulated controls. Parallel non-washout conditions were included for direct comparison.

Supernatants were collected 24 h after restimulation, and cytokine production (IL-2 and IFN-γ) was quantified by ELISA (STEMCELL Technologies, Catalog# 02006 and 02003). Functional recovery after washout was expressed as cytokine production relative to stimulated controls. Experiments were performed using PBMCs from 10 independent healthy donors. Statistical analysis was performed using donor-matched comparisons with one-way repeated-measures ANOVA followed by multiple-comparison testing.

### Patient-derived PBMC experiments

PBMCs from patients with ulcerative colitis (n = 5 independent donors; BioIVT, Catalog# HUMANPBMC-0002203) were processed and cultured under identical conditions as healthy donor PBMCs. Cells were obtained from commercial vendors and were provided to investigators in a de-identified manner. No identifiable donor information was available to investigators. Cells were stimulated with anti-CD3/CD28 in the presence of increasing concentrations of CP8. Cytokine production was quantified by ELISA after 24 h.

### Murine CD28 cross-reactivity assessment

To evaluate functional cross-reactivity of CP8 with murine CD28, primary splenocytes were isolated from 8-10-week-old C57BL/6 mice. Spleens were mechanically dissociated through a 70-µm cell strainer to obtain single-cell suspensions. Red blood cells were lysed using ammonium-chloride-potassium (ACK) lysis buffer, and cells were washed twice with RPMI 1640 medium supplemented with 10% FBS, L-glutamine, and penicillin-streptomycin (100 U/mL and 100 µg/mL, respectively).

Splenocytes were seeded at 2 × 10^5^ cells per well in 96-well plates and stimulated with plate-bound anti-mouse CD3 (2 µg/mL) and soluble anti-mouse CD28 (1 µg/mL) antibodies. CP8 was added at indicated concentrations (0.005-5 µM) at the time of stimulation. Cells were incubated at 37 °C in 5% CO_2_ for 24 h.

Supernatants were collected and murine IL-2 and IFN-γ concentrations were quantified using ELISA kits (STEMCELL Technologies, Catalog# 02022 and 02020) according to the manufacturer’s instructions. Experiments were performed using splenocytes from independent mice (n = 4 biological replicates), each measured in technical triplicate. Data were analyzed using one-way ANOVA with appropriate post hoc testing.

### Plasma Stability and Microsomal Stability

CP8 stability in mouse plasma was evaluated by incubating the peptide (1 µM final concentration) in pooled mouse plasma at 37 °C. Aliquots were collected at 0, 0.5, 1, 2, 4, and 6 h and quenched with ice-cold acetonitrile containing an internal standard to precipitate plasma proteins. Samples were centrifuged at 15,000 × g for 10 min, and supernatants were analyzed by LC-MS/MS. The percentage of remaining parent compound was calculated relative to time zero. Experiments were performed in triplicate.

Metabolic stability was assessed using pooled mouse and human liver microsomes (ThermoFisher Scientific, Catalog# MSMCPL and HMMCPL). CP8 (1 µM) was incubated with microsomes (0.5 mg/mL protein) in potassium phosphate buffer (pH 7.4) at 37 °C in the presence of an NADPH-regenerating system. Reactions were initiated by addition of NADPH and terminated at defined time points (0, 5, 15, 30, 45, and 60 min) by addition of ice-cold acetonitrile.

Samples were centrifuged, and supernatants were analyzed by LC-MS/MS. The natural logarithm of percentage remaining compound was plotted versus time to determine the first-order elimination rate constant (k). Intrinsic clearance (Cl_int_) and microsomal half-life (t_1/2_) were calculated using standard equations. Experiments were performed in triplicate.

### Animal studies

Mice were housed and handled according to the guidelines approved by the Institutional Animal Care and Use Committee (IACUC) of our institution (protocol 2023-0028).

### PK analysis of CP8 in mice

PK studies were conducted in male C57BL/6 mice (8-10 weeks old; 20-25 g). Animals were housed under standard conditions with ad libitum access to food and water. CP8 was formulated in sterile PBS containing 5% DMSO and administered via subcutaneous (s.c.) injection at doses of 1 mg/kg or 5 mg/kg (n = 4 per dose group). Blood samples (∼50-75 µL) were collected via submandibular venipuncture at the following time points post-dose: 0.25, 0.5, 1, 2, 4, 8, 12, and 24 h. Blood was collected into EDTA-coated tubes and centrifuged at 3,000 × g for 10 min at 4 °C to obtain plasma. Plasma samples were stored at −80 °C until analysis.

Plasma concentrations of CP8 were quantified using a validated liquid chromatography-tandem mass spectrometry (LC-MS/MS) method. Briefly, plasma proteins were precipitated using acetonitrile containing an internal standard, followed by centrifugation and injection of the supernatant onto a reverse-phase C18 analytical column. Chromatographic separation was achieved using a gradient of water and acetonitrile containing 0.1% formic acid. Detection was performed in positive electrospray ionization mode using multiple reaction monitoring (MRM). Calibration curves were generated using spiked plasma standards over a concentration range of 1-10,000 ng/mL. The lower limit of quantification (LLOQ) was 1 ng/mL. Quality control samples at low, medium, and high concentrations were included in each analytical run to ensure accuracy and precision.

PK parameters, including maximum plasma concentration (C_max_), time to maximum concentration (T_max_), area under the plasma concentration-time curve from 0 to 24 h (AUC_0-24_ h), terminal elimination half-life (t_1/2_), and apparent clearance (CL/F), were calculated using non-compartmental analysis in Phoenix WinNonlin (Certara). Terminal half-life was determined from the slope of the log-linear terminal phase. Data are reported as mean ± SD.

### Adoptive T Cell Transfer Model of Chronic Colitis

Chronic colitis was induced using the adoptive CD4⁺CD45RB^high^ T-cell transfer model as previously described (34–36). Briefly, spleens were harvested from 5-week-old BALB/c donor mice (Jackson Laboratory), and single-cell suspensions were prepared by mechanical dissociation followed by filtration through a 70-µm nylon mesh. CD4⁺ T cells were enriched by magnetic bead separation, and CD4⁺CD45RB^high^ T cells were purified by fluorescence-activated cell sorting (FACS). Purified cells were washed twice with sterile phosphate-buffered saline (PBS) and resuspended at 1.5 × 10^6^ cells/mL in cold PBS.

Seven-week-old C.B-17 scid recipient mice (Taconic Biosciences) received 3 × 10^5^ CD4⁺CD45RB^high^ T cells by intravenous injection (200 µL per mouse). Mice were randomized into treatment groups (n = 8 per group) following cell transfer.

CP8 was formulated in sterile PBS containing 5% DMSO and administered by subcutaneous injection at 5 mg/kg using either a daily dosing regimen or an intermittent dosing regimen (every other day). Acazicolcept was included as a comparator biologic and administered according to an intermittent dosing schedule (1 mg/kg, twice weekly). Vehicle-treated mice received formulation buffer alone. Treatment was initiated on day 7 post-transfer and continued through day 30.

Disease progression was monitored longitudinally using a disease activity index (DAI) incorporating body weight loss, stool consistency, and fecal blood. Body weight was recorded three times per week. Stool consistency was scored as 0 (normal), 1 (soft but formed), 2 (loose), and 3 (diarrhea). Fecal blood was assessed using a guaiac-based assay and scored as 0 (negative), 1 (trace), 2 (positive), and 3 (gross bleeding). The DAI was calculated as the composite score of these parameters.

At the study endpoint (day 30), mice were euthanized and colons were excised and measured from cecum to rectum under standardized tension. Blood was collected via cardiac puncture, and serum was isolated by centrifugation at 3,000 × g for 10 min at 4 °C. Serum TNF-α and IL-6 levels were quantified using commercial ELISA kits (Thermo Fisher Scientific, Catalog# BMS607-3 and KMC0061) according to the manufacturers’ instructions.

### Statistical analysis

Data are presented as mean ± SEM unless otherwise indicated. Statistical analyses were performed using GraphPad Prism 10.4.1 software. Comparisons between groups were conducted using one-way or two-way analysis of variance (ANOVA) with appropriate post hoc tests as specified in figure legends. *P* values less than 0.05 were considered statistically significant.

## Author contributions

The manuscript was written through contributions of all authors. All authors have given approval to the final version of the manuscript. Conceptualization: M.T.G. and H.D. Methodology: K.K., S.U., and Y.G. Investigation: K.K., S.U., and Y.G. Formal analysis: K.K., S.U., and Y.G. Visualization: K.K., S.U., and Y.G. Supervision: M.T.G. and H.D. Writing-original draft: K.K., S.U., Y.G., M.T.G., and H.D. Writing-review and editing: M.T.G. Funding acquisition: M.T.G. and H.D. Data curation: K.K., S.U., and Y.G. Validation: K.K., S.U., and Y.G. Project administration: M.T.G.

## Data and materials availability

All data needed to evaluate the conclusions in the paper are present in the paper and/or the Supplementary Materials.

