## Supporting Information for "Reversible CD28 checkpoint modulation by cyclic peptides outperforms biologic blockade under exposure-limited conditions"

*Electronic Supplementary Information for*

### Contents

Phage ELISA analysis of selected clones S3
Microscale thermophoresis analysis of CP1 binding to CD28 S4
Microscale thermophoresis analysis of CP2 binding to CD28 S4
Cyclic peptides inhibit CD28-dependent costimulatory signaling in cells S5
CP8 exhibits comparable initial inhibitory efficacy to clinically advanced CD28-targeting biologics in patient-derived PBMCs S6
CP8 functionally inhibits murine CD28-mediated T-cell activation S7
Mass spectrum of CP1 S8
Mass spectrum of CP2 S9
Mass spectrum of CP3 S10
Mass spectrum of CP5 S11
Mass spectrum of CP6 S12
Mass spectrum of CP7 S13
Mass spectrum of CP8 S14

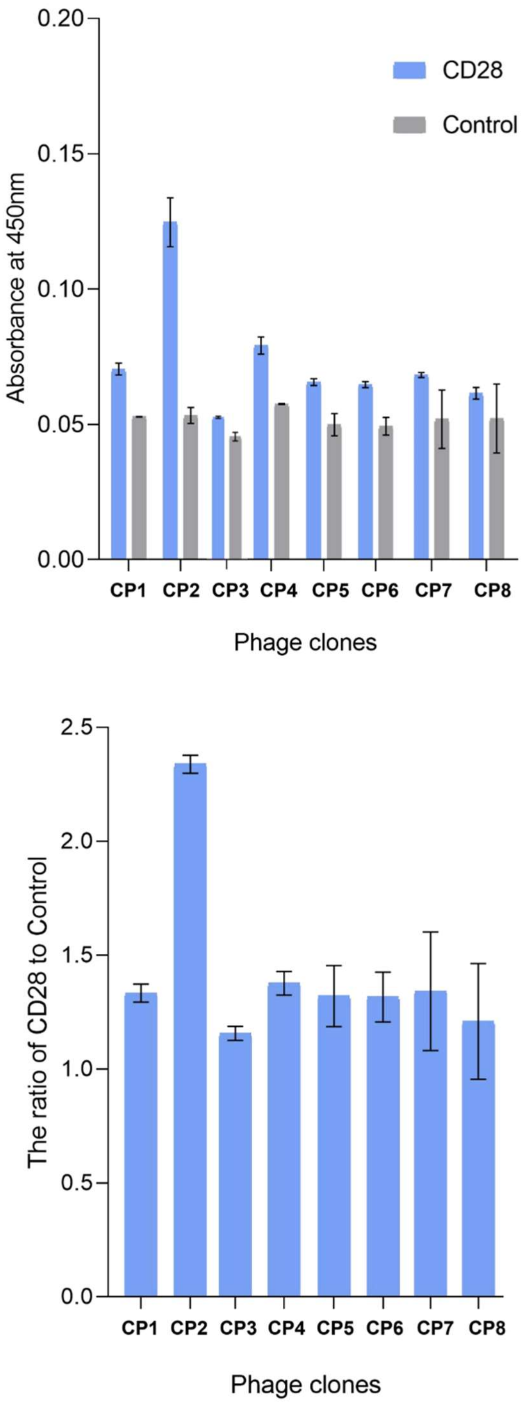


**Figure S1.** Phage ELISA analysis of selected clones. The upper panel displays the raw absorbance values measured at 450 nm, while the lower panel presents the signal ratio of CD28 binding relative to the empty control (E/C), indicating the specificity of peptide binding to CD28. Control wells were prepared without protein coating, containing only PBS buffer.

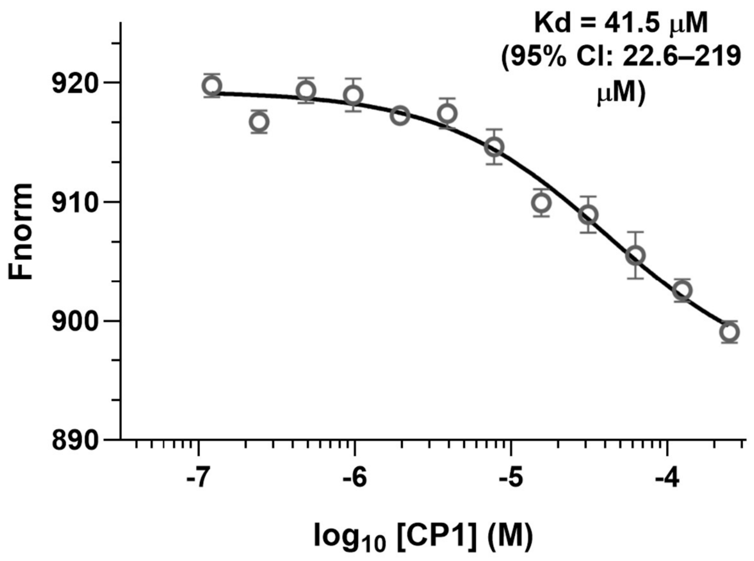


**Figure S2. Microscale thermophoresis analysis of CP1 binding to CD28.** Binding was quantified by MST using intrinsic fluorescence detection (spectral shift mode; Monolith X). Fnorm, (670/650 nm) is plotted as a function of peptide concentration. Data points represent mean ± SD from three independent measurements. Binding curves were fitted using nonlinear regression assuming a 1:1 binding model to estimate equilibrium dissociation constants (Kd).

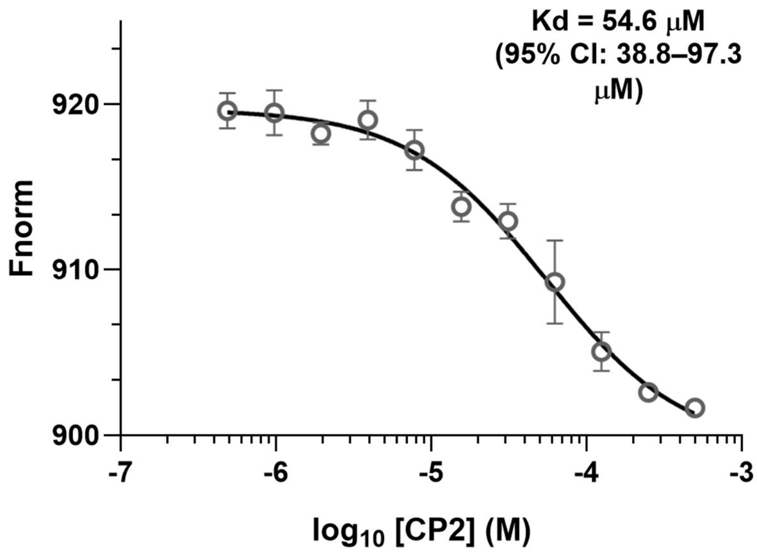


**Figure S3. Microscale thermophoresis analysis of CP2 binding to CD28.** Binding was quantified by MST using intrinsic fluorescence detection (spectral shift mode; Monolith X). Fnorm, (670/650 nm) is plotted as a function of peptide concentration. Data points represent mean ± SD from three independent measurements. Binding curves were fitted using nonlinear regression assuming a 1:1 binding model to estimate equilibrium dissociation constants (Kd).

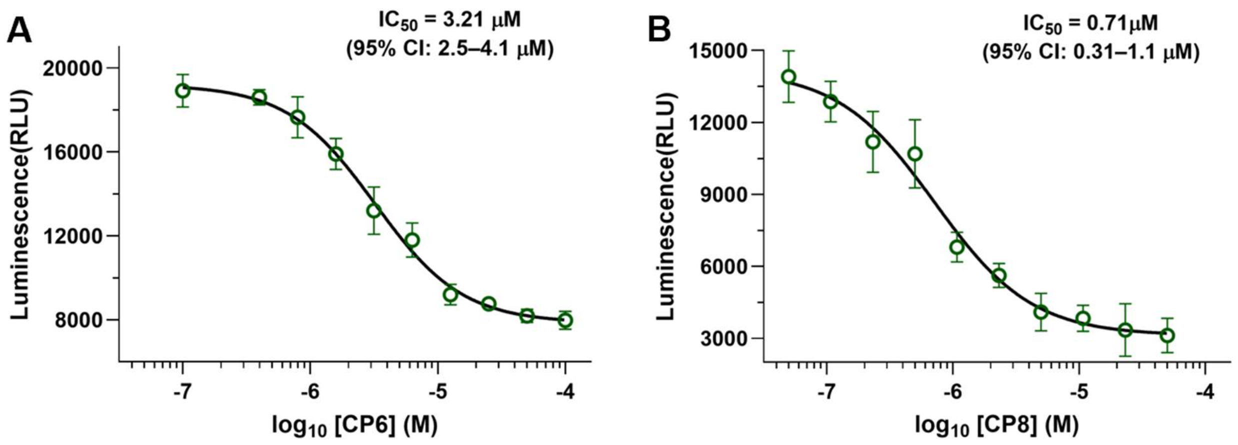


**Figure S4. Cyclic peptides inhibit CD28-dependent costimulatory signaling in cells.** Functional antagonism of CD28 signaling was evaluated using a reporter assay in which Jurkat T cells were co-cultured with antigen-presenting cells to induce CD28-dependent activation, quantified by an NFAT-driven luciferase readout. **(A)** CP6 inhibits CD28 signaling in a concentration-dependent manner with moderate potency. **(B)** CP8 exhibits substantially enhanced inhibitory activity, achieving submicromolar inhibition. Data represent mean ± SEM from at least three independent experiments. Concentration-response curves were fitted using a four-parameter logistic regression model.

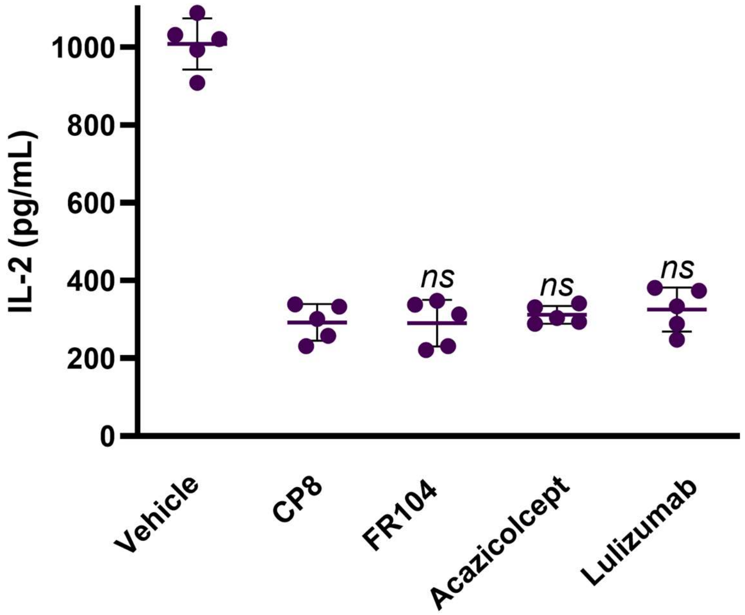


**Figure S5. CP8 exhibits comparable initial inhibitory efficacy to clinically advanced CD28-targeting biologics in patient-derived PBMCs.** PBMCs from patients with ulcerative colitis were stimulated with anti-CD3/CD28 in the presence of CP8, Pegrizeprument (FR104), Acazicolcept, or Lulizumab at 500 nM. All agents produced comparable suppression of IL-2, with no statistically significant differences relative to CP8 (ns). Data are shown as individual donor values with mean ± SEM.

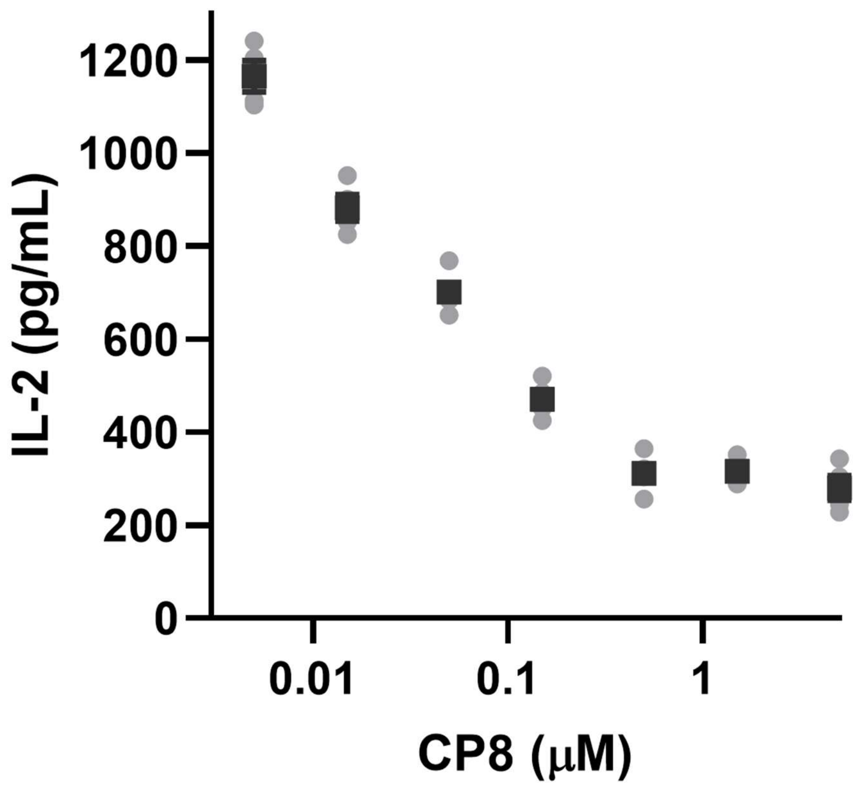


**Figure S6. CP8 functionally inhibits murine CD28-mediated T-cell activation.** Primary splenocytes isolated from C57BL/6 mice were stimulated with anti-CD3 and anti-CD28 antibodies in the presence of increasing concentrations of CP8 (0.03-5 μM). IL-2 production was quantified after 24 h by ELISA. CP8 suppressed murine CD28-dependent T-cell activation in a dose-dependent manner, with an apparent $IC_{50}$ in the nanomolar range comparable to that observed in human PBMC assays. Anti-CD3 stimulation alone produced minimal IL-2 secretion, confirming CD28-dependent costimulatory signaling. Data represent mean ± SEM from independent spleen preparations (n = 4). Statistical analysis was performed using one-way ANOVA with multiple comparisons relative to stimulated control.

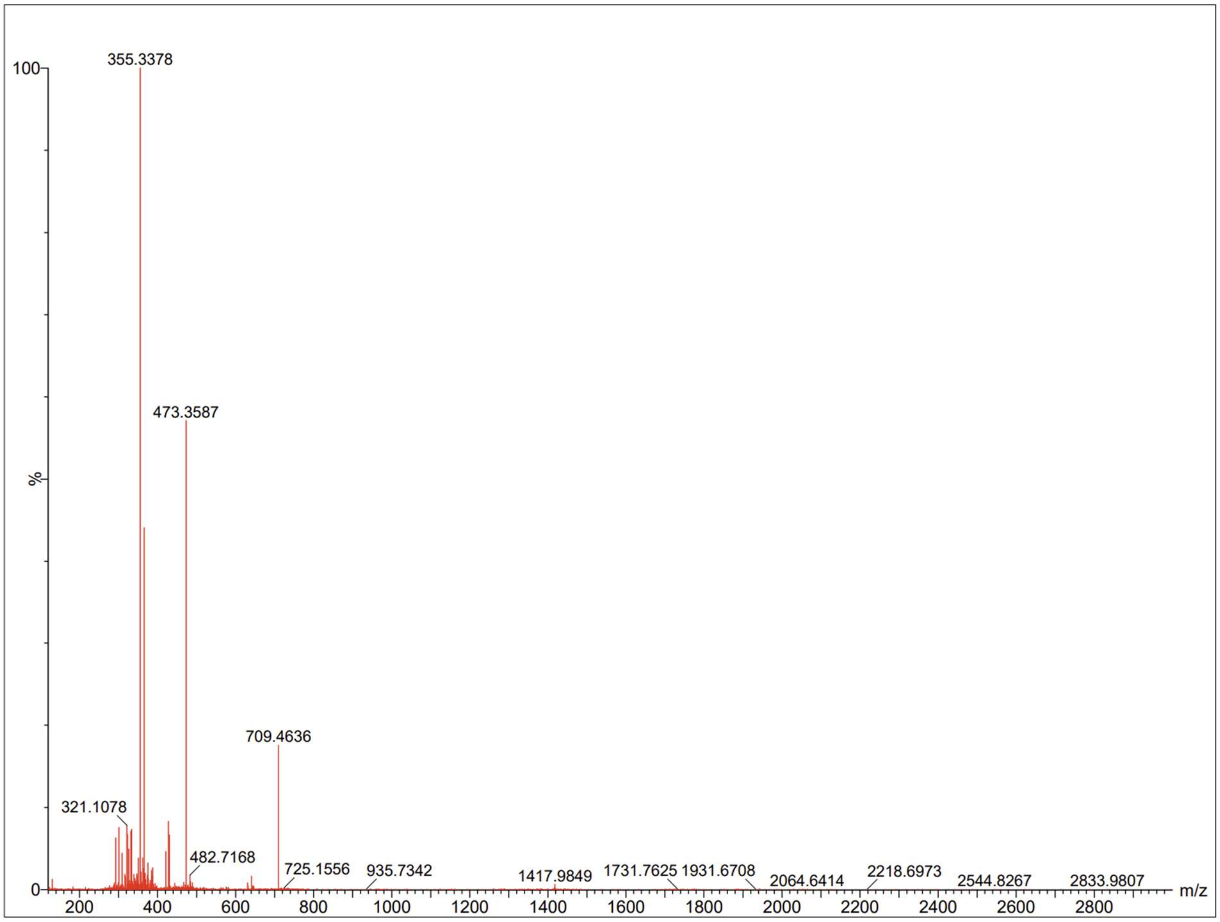


**Figure S7.** Mass spectrum of CP1.

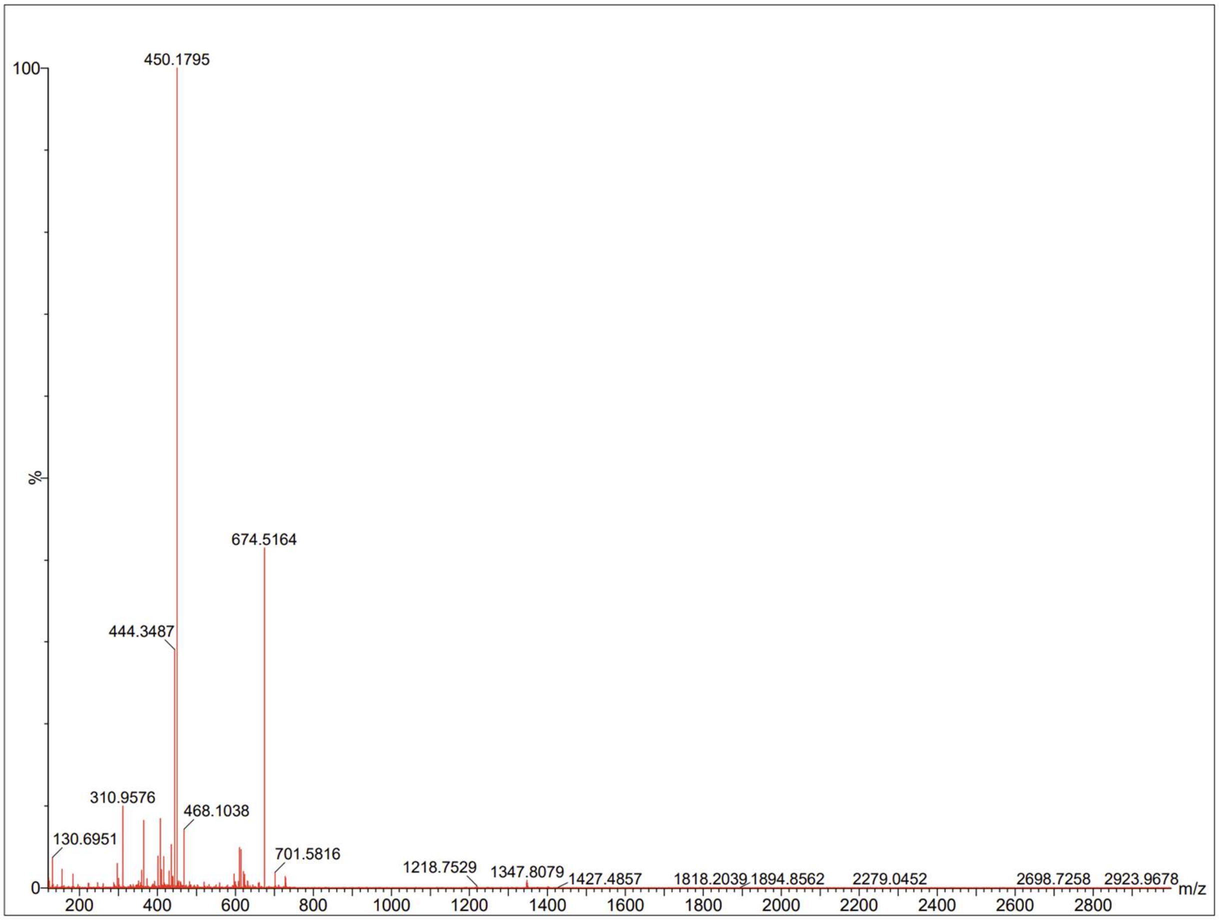


**Figure S8.** Mass spectrum of CP2.

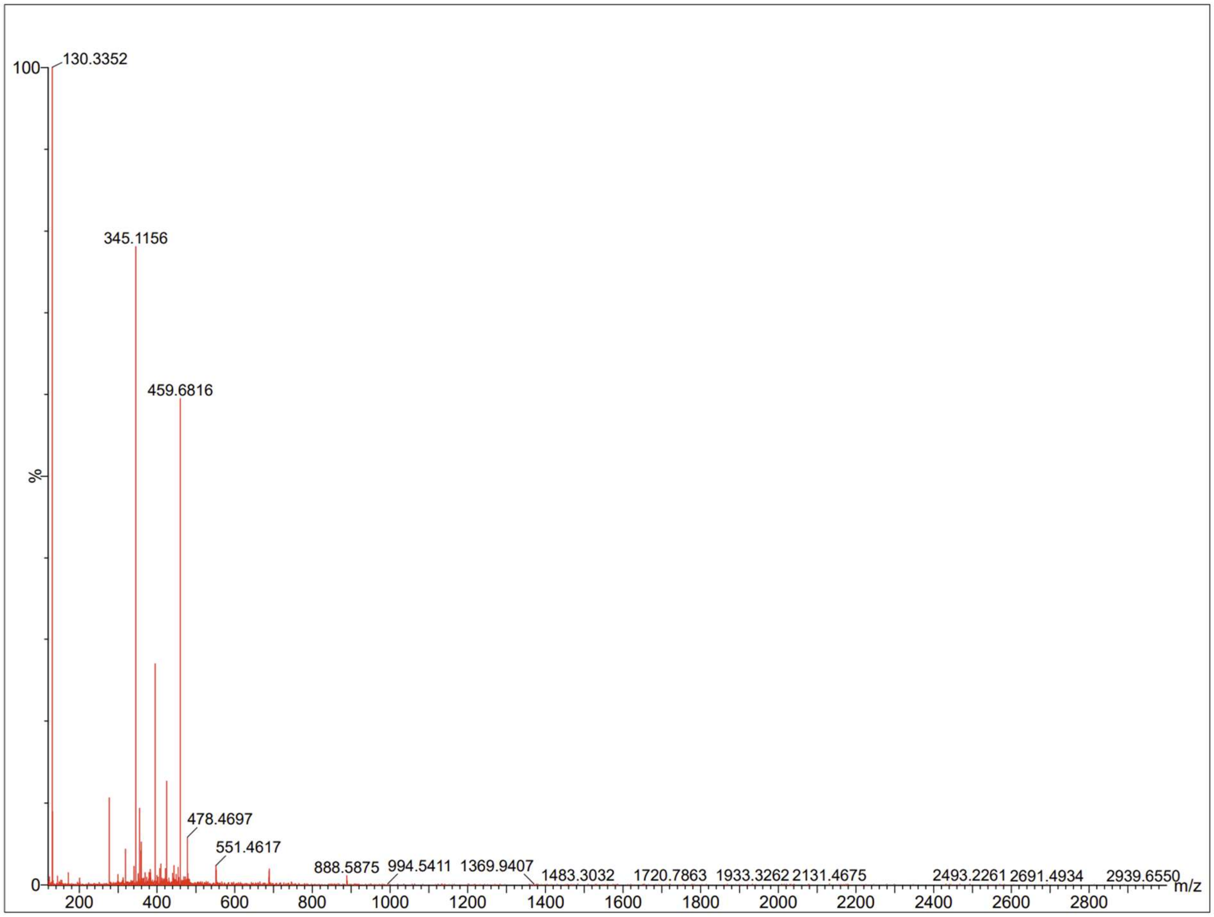


**Figure S9.** Mass spectrum of CP3.

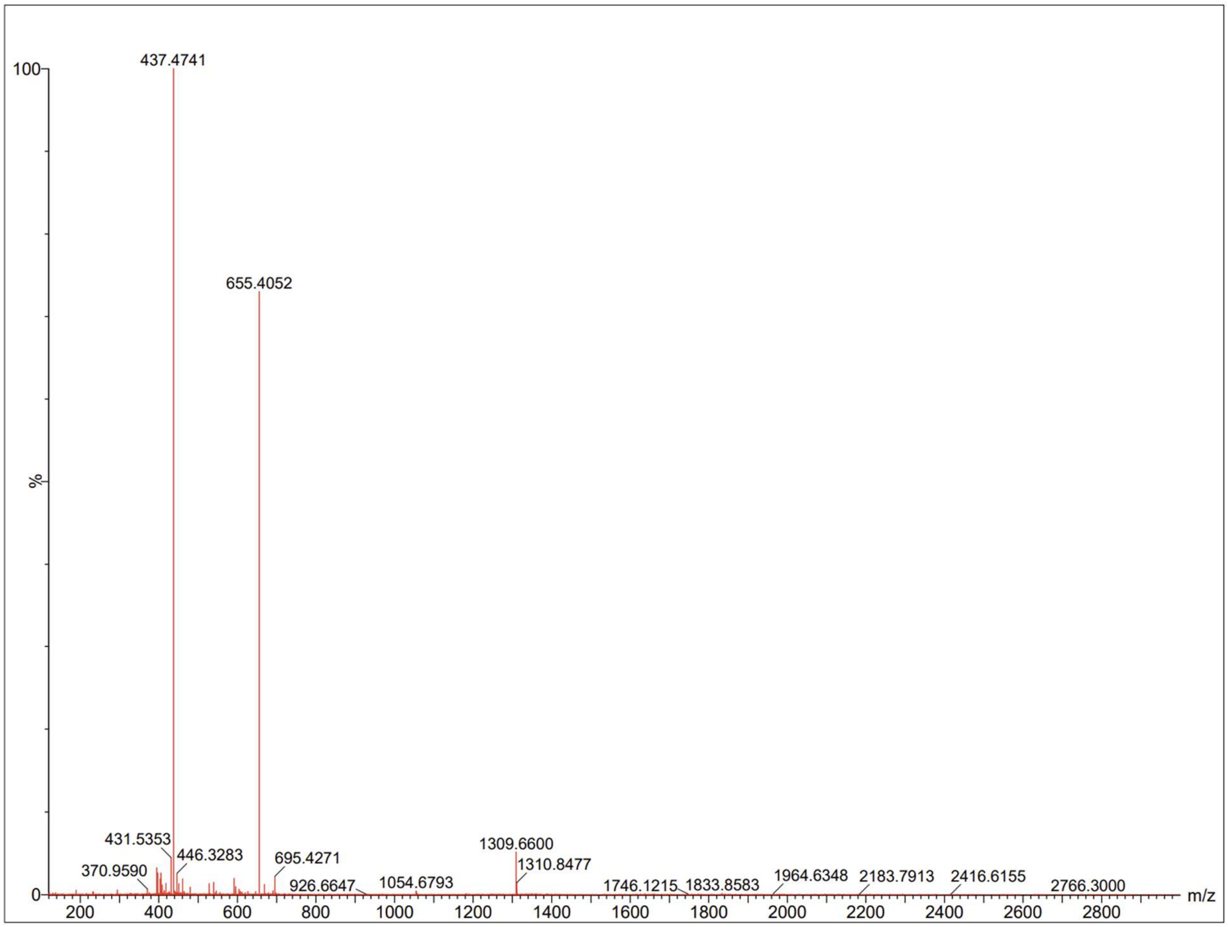


**Figure S10.** Mass spectrum of CP5.

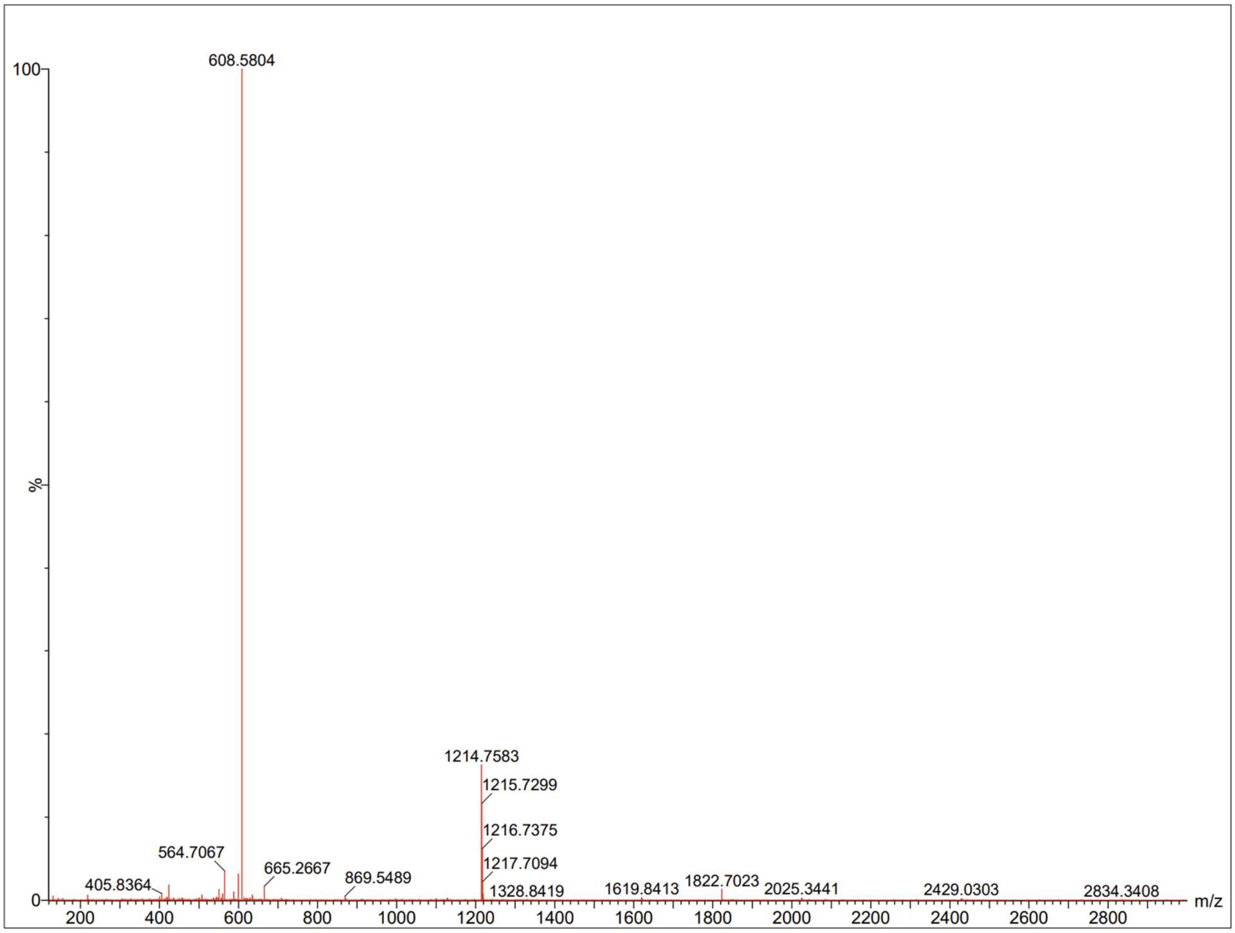


**Figure S11.** Mass spectrum of CP6.

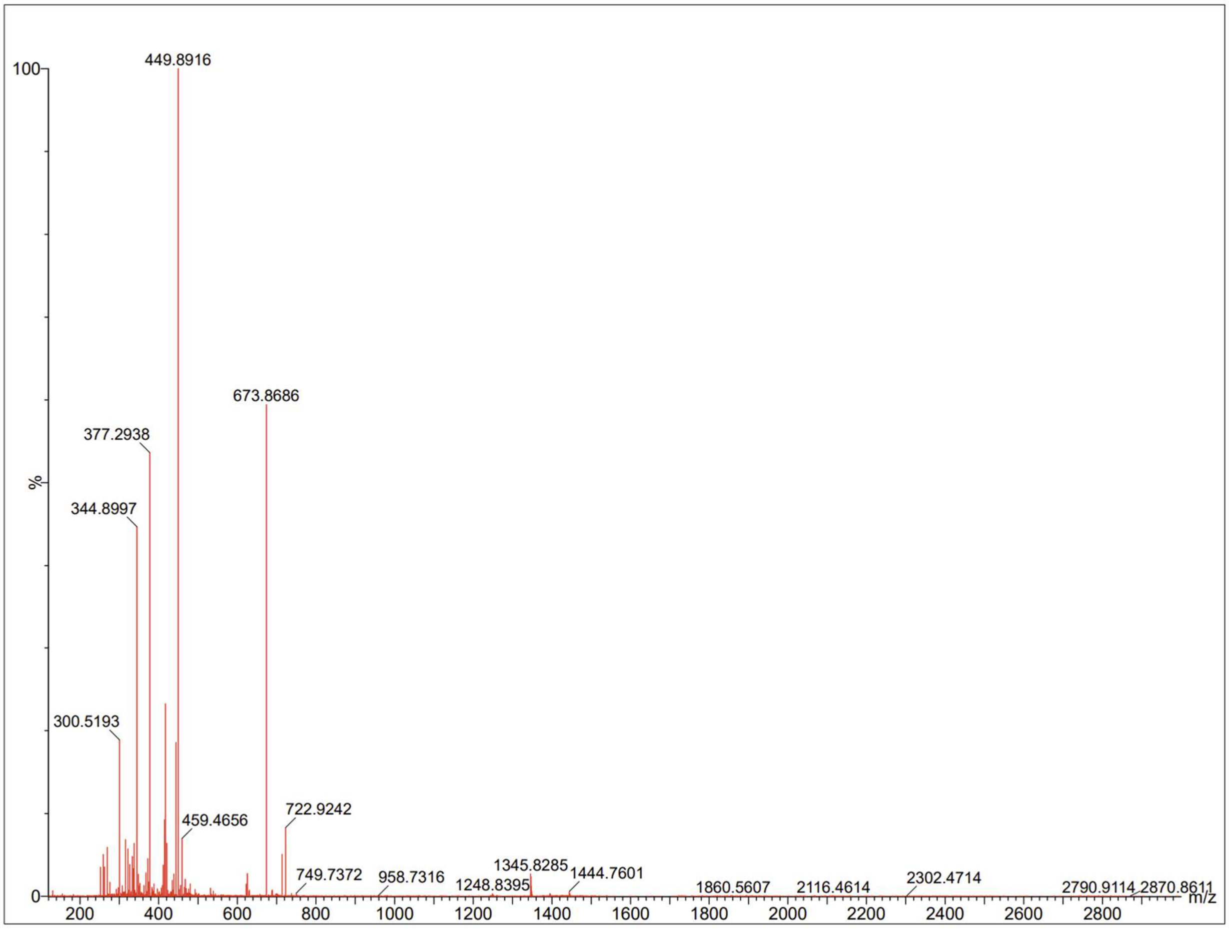


**Figure S12.** Mass spectrum of CP7.

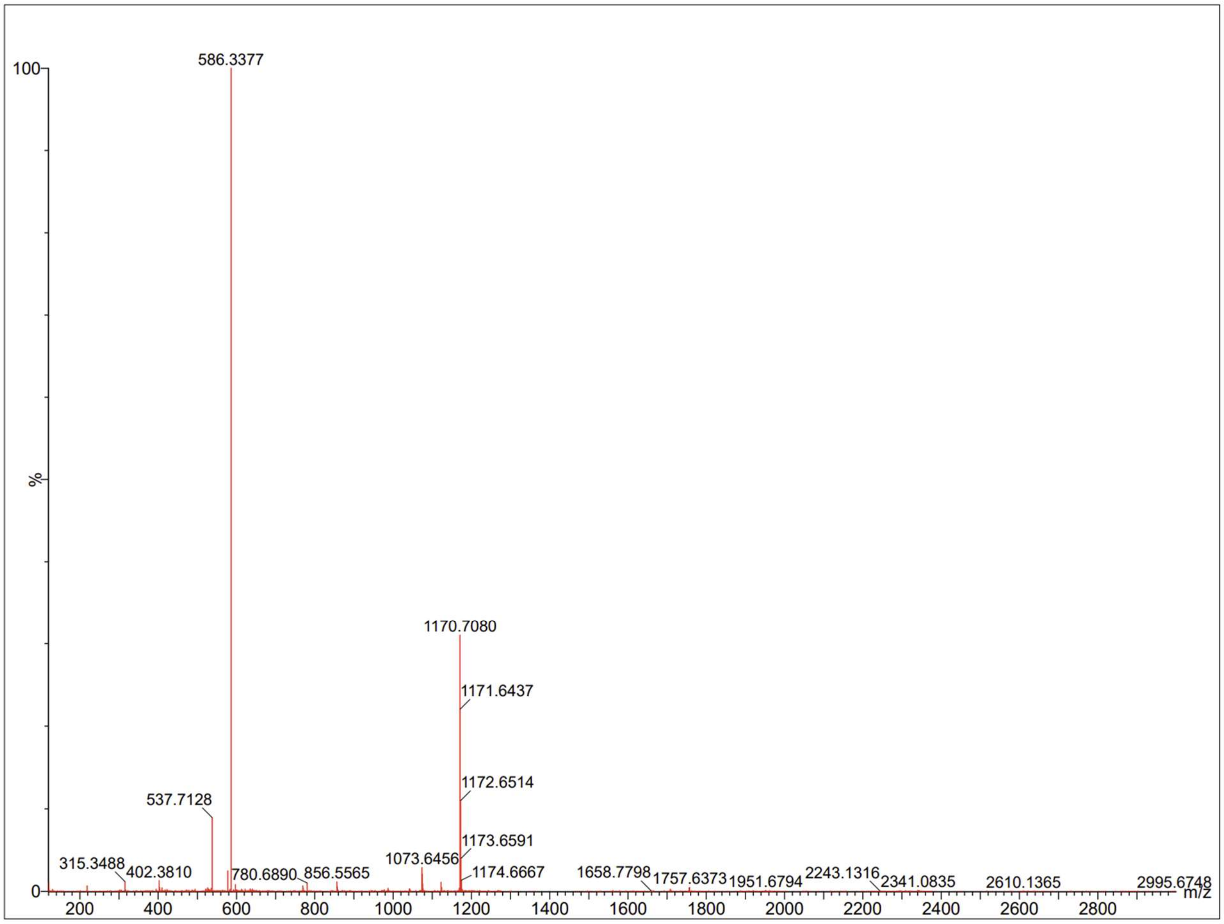


**Figure S13.** Mass spectrum of CP8.
